# Atlas of the Neuromuscular System in the Trachymedusa *Aglantha digitale*: Insights from the advanced hydrozoan

**DOI:** 10.1101/772418

**Authors:** Tigran P. Norekian, Leonid L. Moroz

**Author notes:** Corresponding author: Dr. Leonid L. Moroz, The Whitney Laboratory; University of Florida; 9505 Ocean Shore Blvd. St. Augustine, FL 32080, USA.

## Abstract

Cnidaria is the sister taxon to bilaterian animals, and therefore, represents a key reference lineage to understand early origins and evolution of the neural systems. The hydromedusa *Aglantha digitale* is arguably the best electrophysiologically studied jellyfish because of its system of giant axons and unique fast swimming/escape behaviors. Here, using a combination of scanning electron microscopy and immunohistochemistry together with phalloidin labeling, we systematically characterize both neural and muscular systems in *Aglantha*, summarizing and expanding further the previous knowledge on the microscopic neuroanatomy of this crucial reference species. We found that the majority, if not all (∼2500) neurons, that are labeled by FMRFamide antibody are different from those revealed by anti-α-tubulin immunostaining, making these two neuronal markers complementary to each other and, therefore, expanding the diversity of neural elements in *Aglantha* with two distinct neural subsystems. Our data uncovered the complex organization of neural networks forming a functional ‘annulus-type’ central nervous system with three subsets of giant axons, dozen subtypes of neurons, muscles and a variety of receptors fully integrated with epithelial conductive pathways supporting swimming, escape and feeding behaviors. The observed unique adaptations within the *Aglantha* lineage (including giant axons innervating striated muscles) strongly support an extensive and wide-spread parallel evolution of integrative and effector systems across Metazoa.

## **1** INTRODUCTION

The phylum Cnidaria (jellyfishes, polyps, sea anemones, and corals) is one of the five early branching metazoan lineages. Together with Ctenophores and Bilaterians, they were united within a single group of Eumetazoa – the taxon combining all animals with true neural and muscular systems, whereas sponges and placozoans represent the lineages with a primary absence of neurons and muscles (Kozloff, 1990; Brusca and Brusca, 2003; Nielsen, 2012). Recent phylogenomic studies consistently place cnidarians as the sister group to bilaterian animals (Whelan et al., 2015; Arcila et al., 2017; Simion et al., 2017; Whelan et al., 2017). Cnidaria and Bilateria united as the distinct clade of animals with nervous systems, which possible share a common origin. In contrast, ctenophores might evolve neural and muscular systems independently from the common ancestor of Cnidaria+Bilateria (Moroz, 2014; Moroz et al., 2014; Moroz, 2015; Moroz and Kohn, 2016). Thus, Eumetazoa could be a polyphyletic taxon, and Cnidaria is a key reference group to understand the origin of integrative systems in bilaterians. However, cnidarian neuromuscular systems are highly diverse in their structures and forms, which reflect enormous biodiversity within this group (Bullock and Horridge, 1965; Spencer and Arkett, 1984; Mackie, 1990; Satterlie, 2002; 2011; Steinmetz et al., 2012; Satterlie, 2015b; Leitz, 2016).

Most detailed recent studies of cnidarian neuromuscular systems are focused on two reference species (Leitz, 2016): *Hydra* (a highly specialized freshwater hydrozoan, which lost the medusa stage) and *Nematostella* (an anthozoan, with a primary absence of medusas in its life cycle). Unfortunately, Mesdusozoa received less attention of neuroscientists, despite substantial and widespread occurrences of jellyfishes word-wide (see reviews in (Satterlie and Nolen, 2001; Satterlie, 2002; Mackie, 2004; Satterlie, 2011; Steinmetz et al., 2012; Satterlie, 2015b; a)).

Among jellyfishes, there is a distinct family of Rhopalonematidae (Trachylina/Hydrozoa), which completely lost their polyp stages and developed multiple adaptations to (meso)pelagic life. The most accessible species in this family is *Aglantha digitale –* the Northern jellyfish, predominantly living in deep waters (50-200 m). *Aglantha* frequently reaches surfaces due to mixing currents or upwelling in spring-summer time around the San Guam Archipelago (Northwest Pacific).

*Aglantha digitale* is well-known for its fast swimming and escape response and differ from most other jellyfishes in having three subsets of giant axons (motor axons in the bell, ring axon, and tentacle axons) - a perfect example of convergent evolution of conductive systems mediating synchronous coordinating contractions of the body wall and tentacles for jet-like locomotion (Singla, 1978; Roberts and Mackie, 1980; Weber et al., 1982). Surprisingly, the very same giant axons mediate both fast (escape) and slow (fishing) swimming by utilizing two different spikes, with sodium and calcium ionic mechanisms respectively (Mackie and Meech, 1985; Mackie and Meech, 1995b; a; Meech, 2015).

Twelve functional neural and two conductive epithelial systems were described in *Aglantha*, leading to the suggestion that *Aglantha* possesses a functional analog of the central nervous system, in the form of annulus rather than a compact ganglion (summarized by (Mackie, 2004)). *Aglantha* independently (from some bilaterians and ctenophores) evolved striated muscles as an additional adaptation to fast jet-propulsion swimming.

In the past, *Aglantha* has been studied intensively by several researchers both neurophysiologically and structurally (Singla, 1978; Donaldson et al., 1980; Mackie, 1980; Roberts and Mackie, 1980; Weber et al., 1982; Singla, 1983; Kerfoot et al., 1985; Mackie and Meech, 1985; Arkett et al., 1988; Meech and Mackie, 1993b; a; Mackie and Meech, 1995b; a; Meech and Mackie, 1995; Mackie and Meech, 2000; Mackie et al., 2003; Mackie, 2004; Mackie and Meech, 2008). However, little is known about the neurotransmitter organization in *Aglantha*. For example, it was reported that nitric oxide signaling controls swimming, possibly via nitrergic sensory neurons in tentacles (Moroz et al., 2004), and peptidergic innervation of feeding/receptive systems was also proposed (Mackie et al., 1985).

Here, by utilizing scanning electron microscopy (SEM), fluorescent immunocytochemistry for labeling neural elements, and phalloidin for labeling muscle fibers, we present and summarize the organization of the neuromuscular systems in *Aglantha*, which provide a foundation to expand our understanding of this unique system in the future with modern molecular tools.

## 2 MATERIALS AND METHODS

### 2.1 Animals

Adult specimens of *Aglantha digitale* (the family Rhopalonematidae) were collected from the breakwater and held in 1-gallon glass jars in the large tanks with constantly circulating seawater at 10° - 12° C. Experiments were carried out at Friday Harbor Laboratories, the University of Washington in the spring-summer seasons of 2016-2019.

### 2.2 Scanning Electron Microscopy (SEM)

Animals were fixed whole in 2.5% glutaraldehyde in 0.1 M cacodylic-buffered saline (pH=7.6) for 4 hours at room temperature and washed for a few hours in 0.1 M cacodylic acid buffer. Larger animals, more than 1 cm in diameter, were then dissected into a few smaller pieces, while small animals were processed as whole mounts. For secondary fixation, we used 2% osmium tetroxide in 0.1 M cacodylic buffer for 2 hours at room temperature. Tissues were then rinsed several times with distilled water, dehydrated in ethanol (series of 20-minute incubations in 30%, 50%, 70%, 90% and 100% ethanol) and placed in Samdri-790 (Tousimis Research Corporation) for Critical point drying. After the drying process, the tissues were placed on the holding platforms and processed for metal coating on Sputter Coater (SPI Sputter). SEM analyses and photographs were done on the NeoScope JCM-5000 microscope (JEOL Ltd., Tokyo, Japan).

### 2.3 Immunocytochemistry and phalloidin staining

Adult *Aglantha* were fixed overnight in 4% paraformaldehyde in 0.1 M phosphate-buffered saline (PBS) at +4-5° C and washed for several hours in PBS. The fixed animals were then dissected into smaller pieces to get better access to specific organs and areas of interest, and better antibody penetration.

The dissected tissues were pre-incubated overnight in a blocking solution of 6% goat serum in PBS and then incubated for 48 hours at +4-5° C in the primary antibodies diluted in 6% of the goat serum at a final dilution 1:50. The rat monoclonal anti-tubulin antibody (AbD Serotec Cat# MCA77G, RRID: AB_325003) recognizes the alpha subunit of α-tubulin, specifically binding tyrosylated α-tubulin (Wehland and Willingham, 1983; Wehland et al., 1983). Following a series of PBS washes for 8-12 hours, the tissues were incubated for 24 hours in secondary goat anti-rat IgG antibodies: Alexa Fluor 488 conjugated (Molecular Probes, Invitrogen, Cat# A11006, RRID: AB_141373), at a final dilution 1:30 and washed for 12 hours in PBS in refrigerator at +4-5°C.

To label the muscle fibers, we used well-known marker phalloidin (Alexa Fluor 568 phalloidin from Molecular Probes), which binds to F-actin (Wulf et al., 1979). After running the immunocytochemistry protocol, the tissues were incubated in phalloidin solution in PBS for 8 hours at a final dilution 1:80 and then washed in several PBS rinses for 12 hours. The tissues were then mounted on glass microscope slides in Vectashield hard-set mounting medium with DAPI for labeling cell nuclei (Cat# H-1500). The preparations were viewed and photographed using a Nikon research microscope Eclipse E800 with epi-fluorescence using standard TRITC and FITC filters, and Nikon C1 Laser Scanning confocal microscope.

In immunocytochemical double-labeling experiments, we used both anti-α-tubulin antibody and antibody against FMRFamide (rabbit polyclonal; Millipore, Sigma, Cat# AB15348, RRID: AB_805291). After incubating in a blocking solution of 6% goat serum in PBS, the tissues were then placed for 48 hours at +4-5° C in the anti-FMRFamide antibodies diluted in 6% goat serum at a final dilution 1:200. Anti-α-tubulin antibodies were added in the same solution 12 hours after anti-FMRFamide antibodies for the rest of the primary body incubation. After several PBS rinses over 8-12 hours, the tissues were placed for 24 hours in secondary goat anti-rabbit IgG antibodies, TRITC-conjugated (Sigma-Aldrich, Cat# T6778, RRID: AB_261740), at a final dilution 1:60. In 8-10 hours after the start of secondary antibody incubation, goat anti-rat IgG antibodies (Alexa Fluor 488 conjugated) for anti-α-tubulin primary antibodies were added in the same solution for the rest of secondary antibody incubation period. Following 12 hours of washing, the tissue was mounted on glass microscope slides and observed as previously described.

### 2.4 Antibody specificity

Rat monoclonal anti-tyrosinated alpha-tubulin antibody is raised against yeast tubulin, clone YL1/2, isotype IgG2a (Serotec Cat # MCA77G; RRID: AB_325003). The epitope recognized by this antibody has been extensively studied, including details about antibody specificity and relevant assays (Wehland and Willingham, 1983; Wehland et al., 1983). As reported by Wehland et al. (1983) - this rat monoclonal antibody “reacts specifically with the tyrosylated form of brain alpha-tubulin from different species”(Wehland et al., 1983). We have successfully used this specific anti-α-tubulin antibody before on several species of ctenophores to label their nervous system (Norekian and Moroz, 2016; 2019b; c; Norekian and Moroz, 2019a). In addition, we tested the specificity of immunostaining by omitting either the primary or the secondary antibody from the procedure. In both cases, no labeling was detected.

The specificity of anti-FMRFamide antibodies to only FMRFamide has been questioned before, and they have been shown to label an entire family of RFamide neuropeptides, which all are known to be present in a diversity of neural systems across Metazoa (Grimmelikhuijzen and Spencer, 1984; Greenberg et al., 1988). Thus, we used anti-FMRFamide antibody not to locate specifically FMRFamide, but as an additional marker of the neural (peptidergic) elements in *Aglantha*. It worked well in previous studies on *Aglantha* (Mackie et al., 1985), as well as in our current investigation. As a form of control experiments, we omitted the anti-FMRFa antibody or the secondary antibody from the protocol. No labeling was detected in both cases. In addition, when other primary antibodies produced in rabbit replaced anti-FMRFa antibody in the same protocol with the same chemicals, the pattern of labeling was completely different.

## 3 RESULTS

### 3.1 Introduction to the *Aglantha* anatomy

Adult *Aglantha digitale* is usually 1-1.5 cm in length and has a conical transparent bell-like shape with about several dozen reddish tentacles. In the resting or a slow fishing mode, these tentacles are relaxed, but they can rapidly contract during a very fast escape response. This tentacle retraction and whole bell muscle contraction that causes the jet-like propulsion of the animal during escape are mediated by the nervous system with a different subset of axons (Mackie, 2004). Figure 1 provides a broad overview of the *Aglantha* anatomy with the localization of giant axon systems in the subumbrella (the concave inner surface of the bell) and around the margin of the bell.

**Figure 1.**
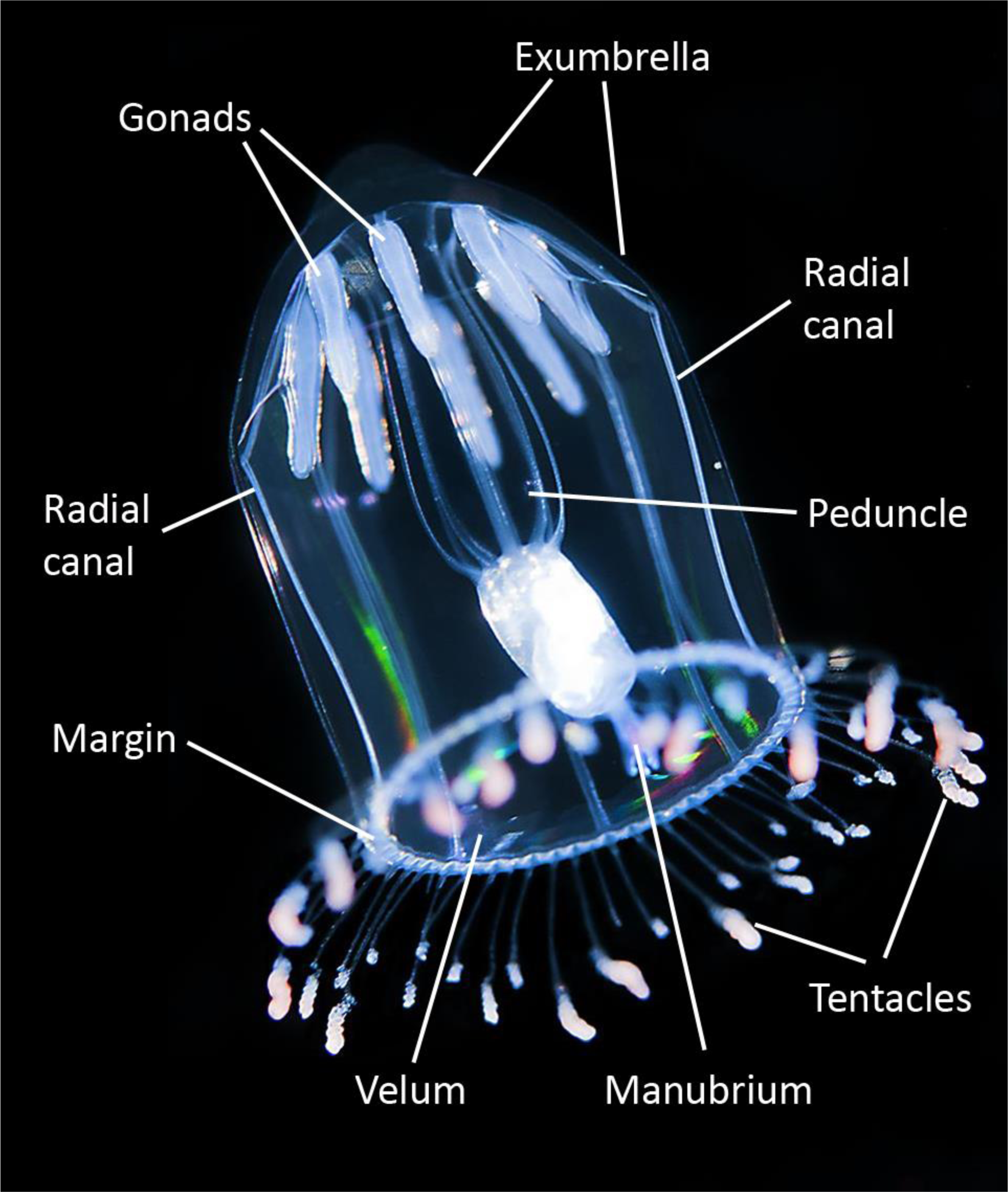
The hydromedusa *Aglantha digitale* and its anatomy. A general view of free-swimming *Aglantha* with major organs visible. The motor giant axons run along the radial canals across the subumbrella surface. The ring nerve with a ring giant axon is located in the margin of the bell. The image of *Aglantha* was taken by Erling Svensen: https://www.artsdatabanken.no/Pages/F21737

Eight statocysts (see details below) are located in the bell-margin areas between the eight radial canals. These statocysts on the umbrella margin of *Aglantha* have been described before and shown to provide a sense of gravity and control direction of swimming (Singla, 1978; Mackie, 1980).

There are eight gonads visible through the bell together with the gastric peduncle and the manubrium carrying the mouth complex, which has four lips in *Aglantha*.

### 3.2 The structure of the umbrella and manubrium

The umbrella wall (Fig. 1) consists of several layers of different cell types. The outside layer of the exumbrella is a layer of epithelial cells 20-30 µm in diameter (Fig. 2a). These cells were not labeled by α-tubulin antibody (AB) or phalloidin. However, phalloidin frequently outlined the borders between these cells, making them visible. On the other side, the external layer of subumbrella is a layer of myoepithelial cells. These cells are elongated in the radial direction with underlying striated muscle tails oriented in the circumferential direction (Singla, 1978; Kerfoot et al., 1985).

**Figure 2.**
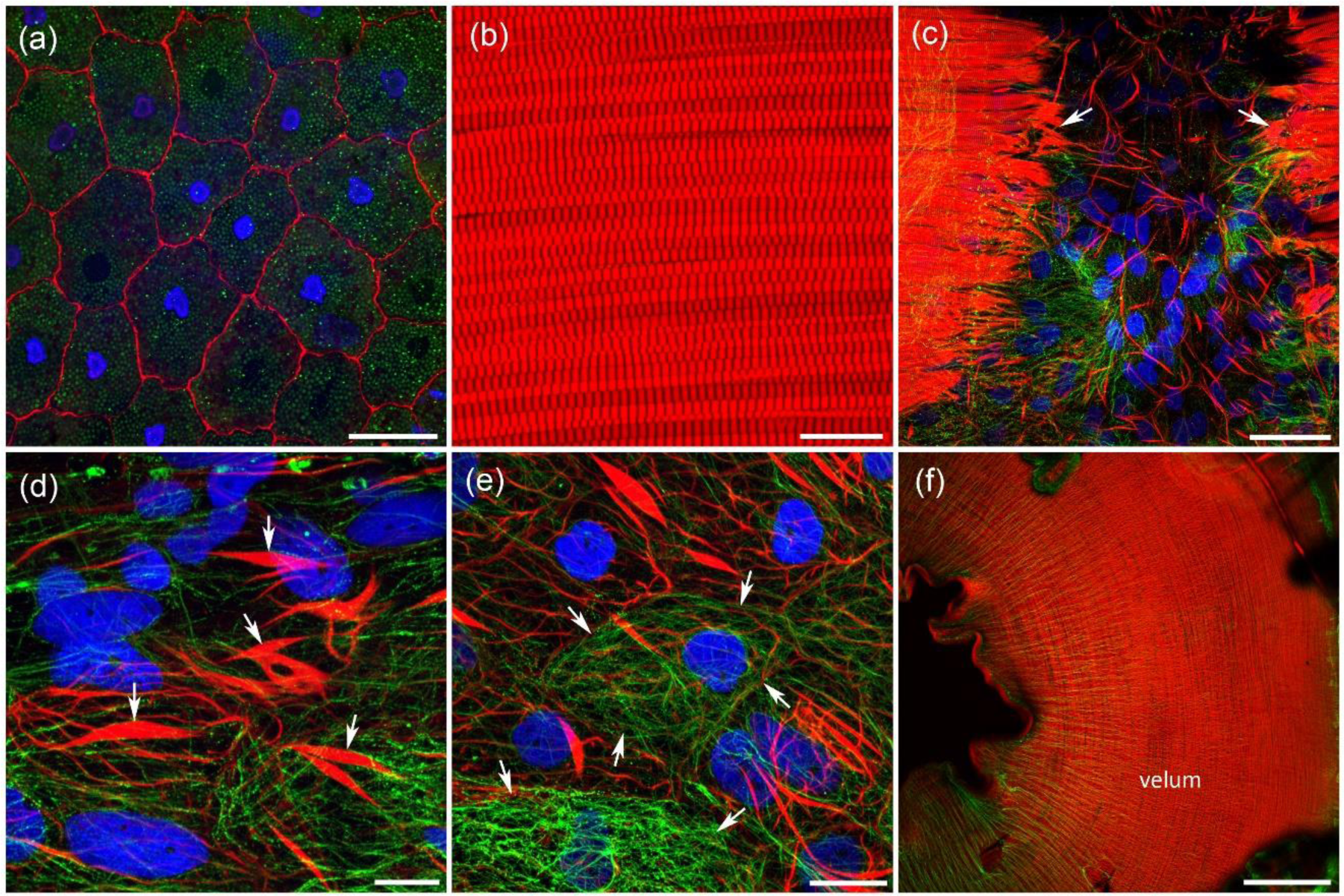
The structure of the umbrella body wall in *Aglantha*. α-Tubulin antibody (AB) is green, phalloidin is red, and nuclear DAPI staining is blue. (**a**) The outside layer of exumbrella is a layer of large epithelial cells, which borders are labeled by phalloidin. (**b**) The layer of circular striated muscles, which powerful contractions cause the propulsion movements of *Aglantha*. These muscles are parts of the myoepithelium and located close to the subumbrella surface. (**c**) The layer of striated muscles is brightly stained by phalloidin and can mask other structures. However, if striated muscles are cut and pulled aside, as indicated by arrows, additional details can be revealed. (**d**) Numerous individual smooth muscle fibers (arrows) in the body wall, which have the shape of a spindle. (**e**) At high magnification individual intracellular filaments, labeled by α-tubulin antibody, could be seen inside the cells (cell borders are indicated by arrows). (**f**) The velum at the bottom of the umbrella with a layer of circular muscles. Scale bars: a - 25 µm; b, d - 10 µm; c - 40 µm; e - 15 µm; f - 250 µm.

The round-shaped and elongated, in a radial direction, cell bodies of these myoepithelial cells were visible in scanning electron microscopy (SEM) images (Fig. 3c). The layer of striated muscle fibers was always brightly labeled by phalloidin (Fig. 2b). The synchronous contraction of striated muscle fibers forces water out of the bell and induces forward movement of *Aglantha*. The intense staining of the striated muscle layer prevented seeing other details in the body wall. However, when we cut and pulled apart the striated muscles, we could observe additional components in the umbrella wall (Fig. 2c), including individual smooth muscle fibers (Fig. 2d,e), which were oriented in different directions. These muscles were presumably responsible for local body movements and body shape changes. Phalloidin also labeled a layer of circular muscles in the velum; their contractions reduced the aperture and, therefore, increasing the force of propulsion (Fig. 2f).

**Figure 3.**
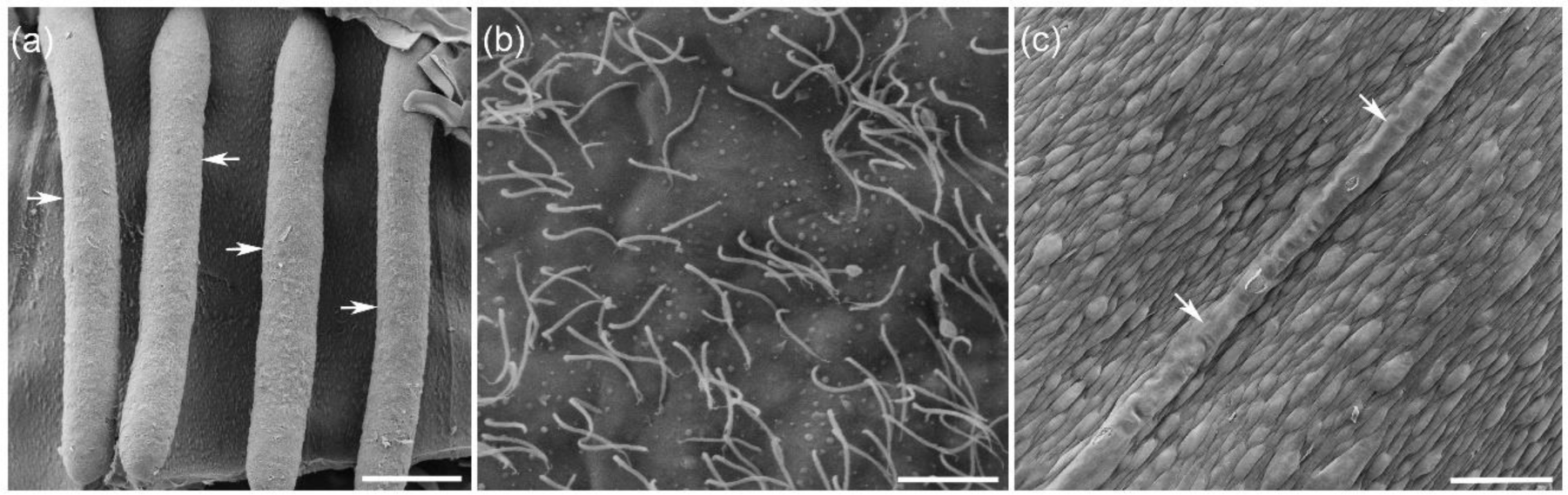
Scanning Electron Microscopy (SEM) of subumbrella surface in *Aglantha*. (**a**) Gonads (arrows) are attached to the subumbrella surface at their upper end. (**b**) The entire surface of the gonads is covered with cilia. (**c**) The subumbrella surface with the canal (arrows) running from the velum to the apex that contains motor giant axon. Elongated round-shaped cell bodies of myoepithelium can be seen across the surface. Scale bars: a - 500 µm; b - 10 µm; c - 100 µm.

The subumbrella region, inside the bell, includes two structures – the gonads and manubrium with the mouth, which also functions as the anus. Eight long cylindrical gonads are evenly spread in a circle around the wall of a bell and attached by one side to subumbrella closer to the apex (Fig. 3a). The surface of gonads is covered by thin cilia (Fig. 3b).

The long manubrium is a tubular structure that has a mouth at its end with four lips or reduced arms (Fig. 4a,b). On the other side, the manubrium is attached to the peduncle and then the subumbrella. The manubrium has muscular walls and always brightly labeled by phalloidin (Fig. 5a,b). The peduncle has much thinner walls with several thin, mostly longitudinally oriented muscle fibers also stained by phalloidin (Fig. 5e). Closer to the subumbrella, the smooth muscle fibers become shorter and spindle-shaped, looking much like those in the wall of the umbrella (Fig. 5f).

**Figure 4.**
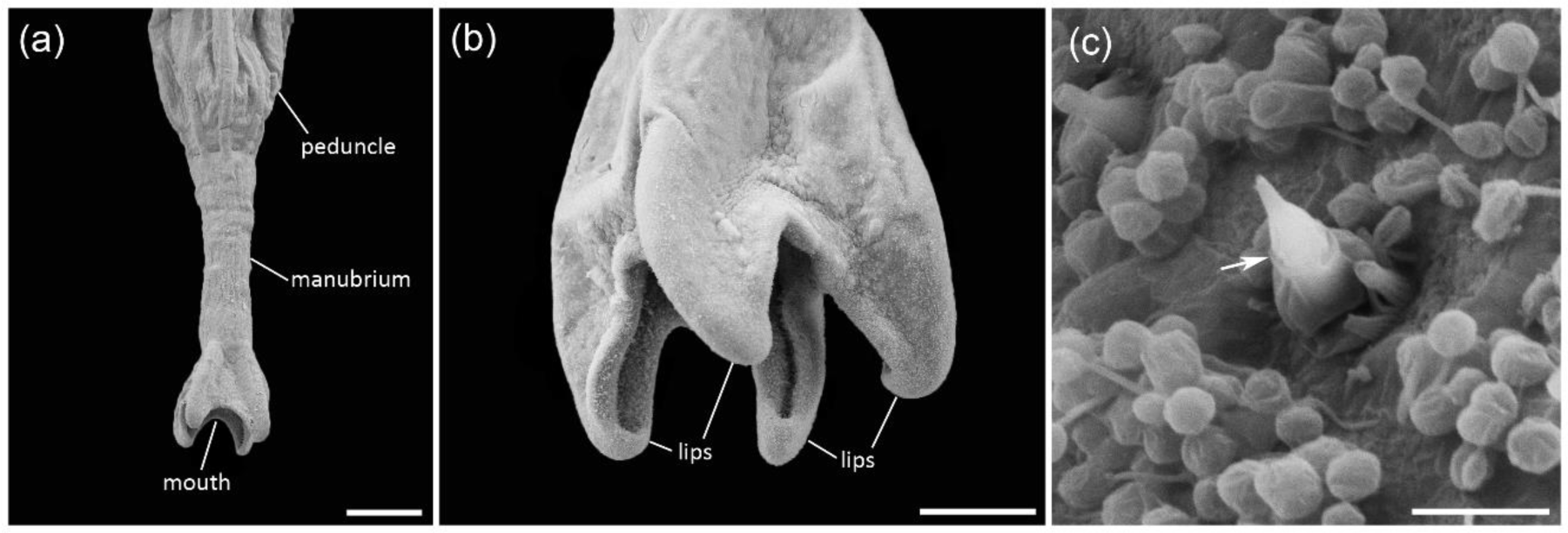
Scanning Electron Microscopy (SEM) of the manubrium and mouth area in *Aglantha*. (**a**) Low magnification shows the manubrium in its entire length attached to the peduncle. (**b**) The mouth on the lower side of manubrium is surrounded by four lips. (**c**) A single cnidocyte (arrow) on the internal surface of the lips at high magnification. Scale bars: a - 500 µm; b - 200 µm; c - 5 µm.

**Figure 5.**
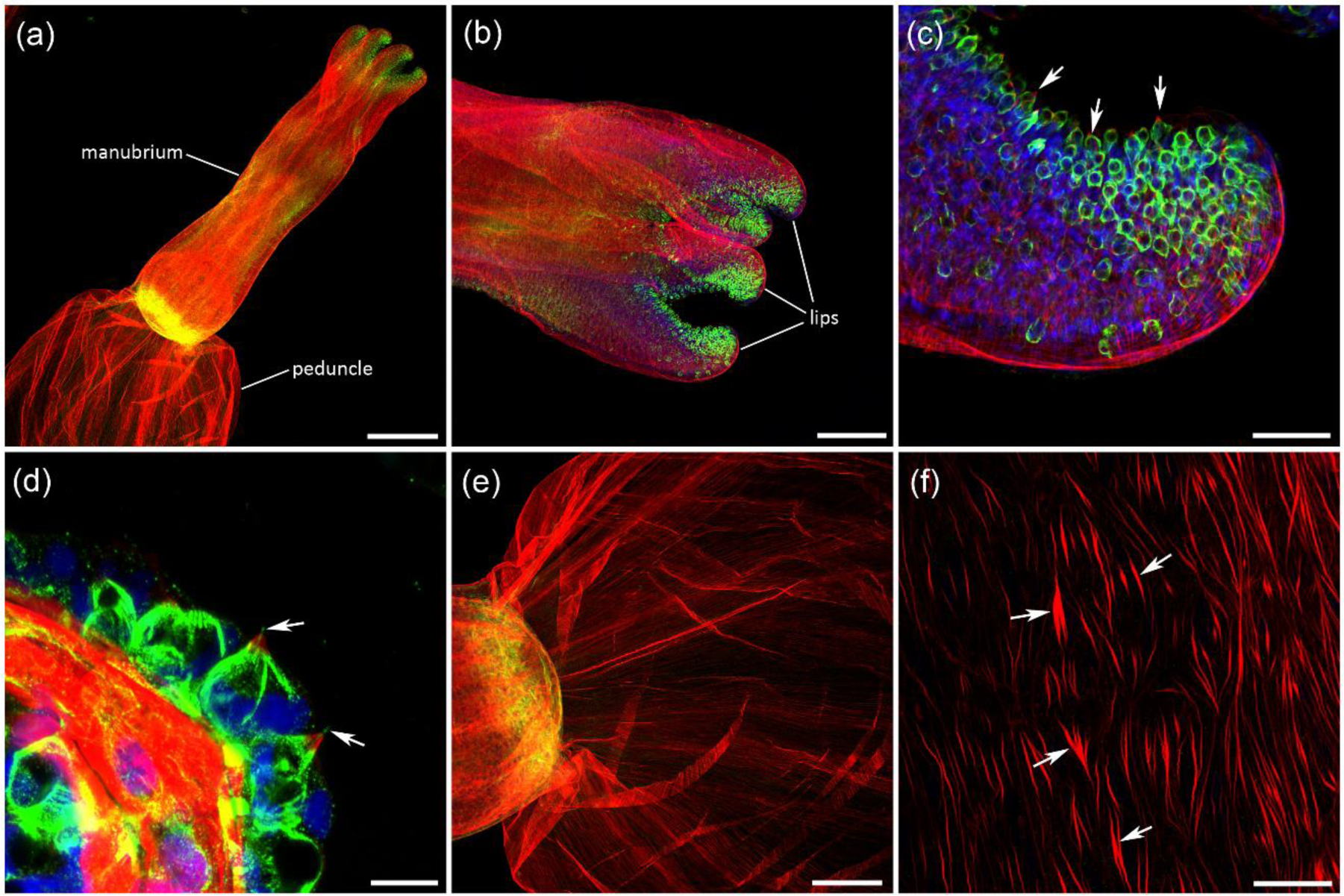
*Aglantha*’s manubrium attached to peduncle on one side, with mouth and four lips on the other side (**a**, **b**). In all images: α-tubulin antibody (AB) labeling is green; phalloidin - red, and DAPI - blue. (**c**) The inner surface of the lips is covered with nematocytes (cnidocytes), which are brightly labeled by α-tubulin AB (arrows). (**d**) Those nematocytes have the same structure as cnidocytes on the tentacles and have a single mechanosensory cilium (cnidocil) labeled by α-tubulin AB (arrows). The surrounding ring of microvilli is labeled by phalloidin (red). (**e**) The thin wall of a peduncle has fine muscle fibers labeled by phalloidin. (**f**) Closer to the umbrella top, the muscle fibers in the peduncle wall become shorter (arrows) and have a spindle-like shape similar to figure 2 (**d**, **e**). Scale bars: a - 500 µm; b, e - 200 µm; c, f - 50 µm; d - 10 µm.

The inner surface of the four lips was covered by nematocytes (or cnidocytes). Nematocytes cell bodies were labeled by α-tubulin AB (Fig. 5b,c). α-Tubulin AB also stained a mechanosensory cilium (cnidocil) that triggers the nematocyst discharge (Fig. 5d). The mechanosensory cilium is surrounded at its base by a ring of microvilli, which are labeled by phalloidin (Fig. 5c,d). The structure of nematocytes on the lips was nearly identical to nematocytes on the tentacles (described later). The presence of nematocytes on the inside surface of the lips was also detected by SEM (Fig. 4c).

The manubrium with its mouth and lips can receive a lot of sensory inputs as well as display fine motor movements during feeding. Therefore, it is not surprising that it has rich innervation. α-tubulin AB labeled a dense network of neural processes in the subepithelial layer of the manubrium wall (Fig. 6a,b). Neural cell bodies, 10-15 µm in diameter, were visible in the nodes of this network at higher magnification (Fig. 6c). The neural network was located above the thick layer of longitudinal and circular muscle fibers (Fig. 6b,c). Neural processes were also found in the lips area, including their inside surface, at the base of nematocytes (Fig. 6d). We have also identified neuron-like cells (10-15 µm in diameter) in the wall of gonads (Fig. 6e,f). Most of these cells were bipolar or tripolar, some producing connections with each other and thus forming a network (Fig. 6e,f).

**Figure 6.**
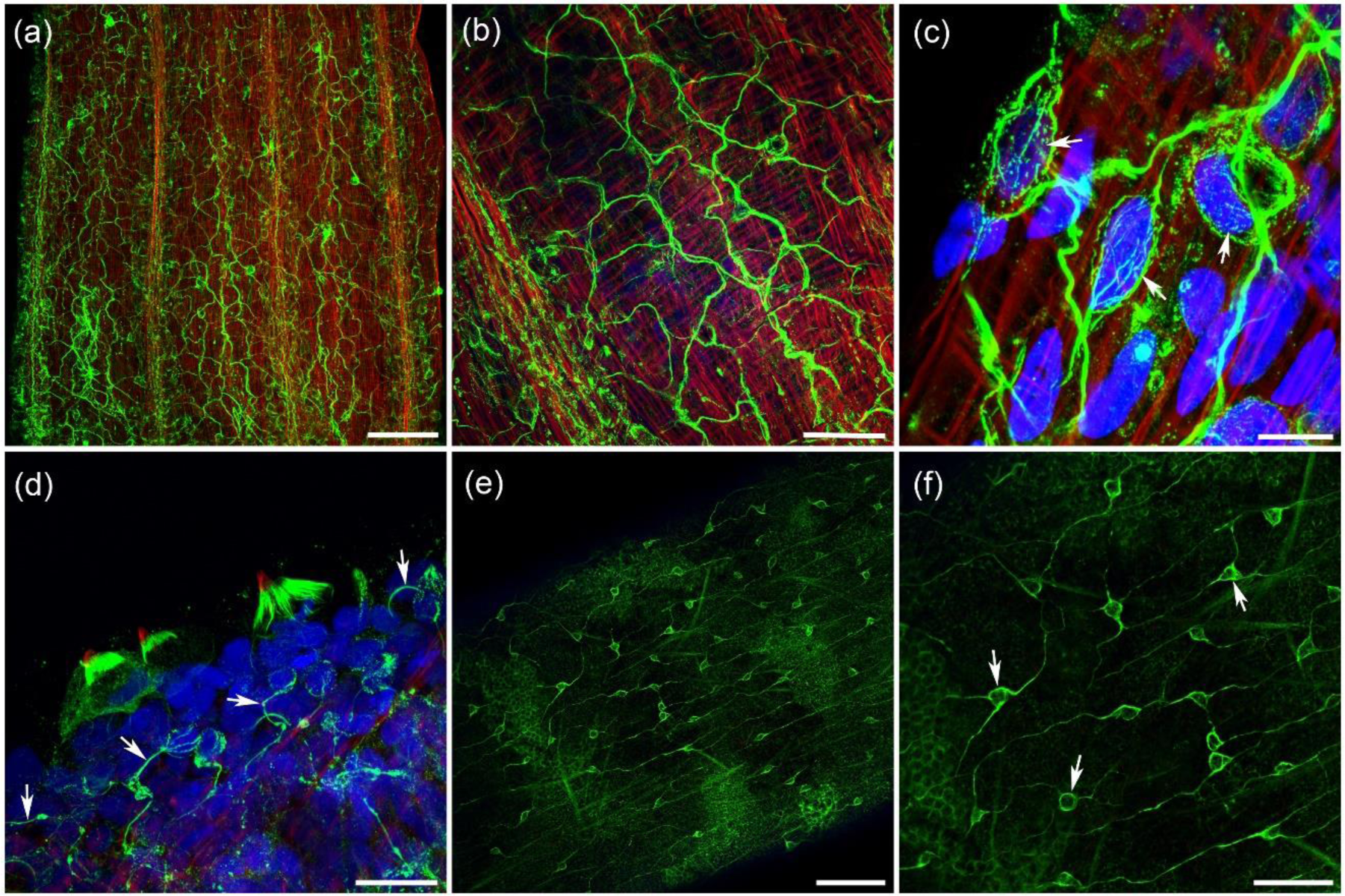
Neural plexus in the manubrium and gonads labeled by α-tubulin AB (green). Phalloidin is red, while nuclear DAPI is blue (**a**, **b**). There is a dense neural plexus in the ectodermal layer of the manubrium in *Aglantha*. (**c**) Higher resolution image shows individual neural cell bodies (arrows) and multiple processes connecting and forming a network-like mesh. (**d**) The nerve processes are next to the nematocytes on the inner surface of the lips (arrows). (**e**, **f**) Neural-like cells labeled by α-tubulin AB form a network on the surface of the gonads (arrows). Scale bars: a, e - 100 µm; b - 40 µm; c - 10 µm; d - 20 µm; f - 50 µm.

### 3.3 Innervation of the umbrella - motor giant axons and nerve rings

The most notable feature, unique to *Aglantha*, is the system of motor giant axons, which play a key role in both slow and fast swimming behavior (Singla, 1978; Donaldson et al., 1980; Roberts and Mackie, 1980; Mackie and Meech, 1985). The motor giant axons were brightly labeled by α-tubulin AB and were easily identifiable in all preparations (Fig. 7a-d). Eight motor giant axons were evenly spread around the umbrella and were running from the umbrella margin to its apex.

**Figure 7.**
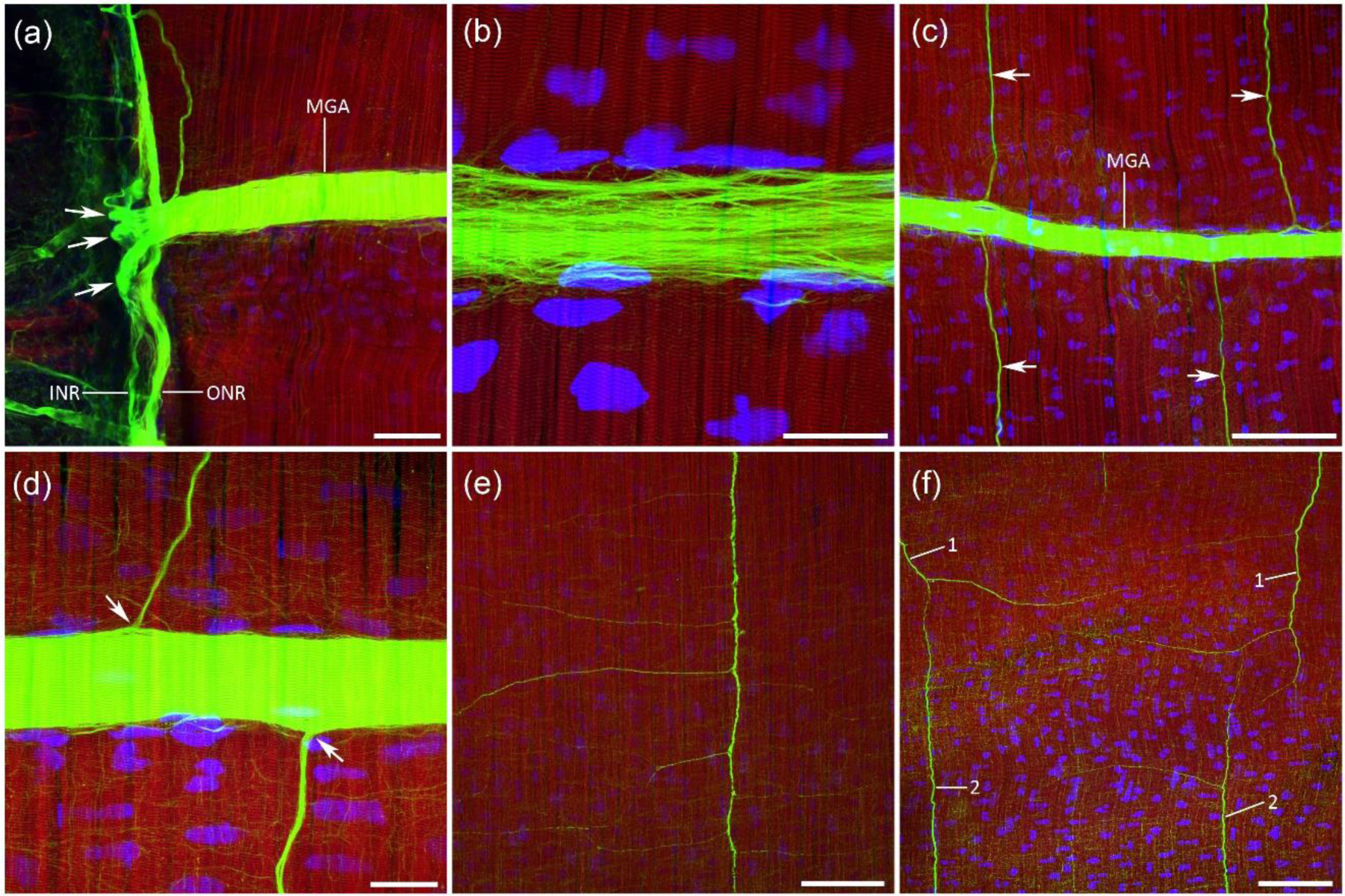
Motor giant axons in *Aglantha* labeled by α-tubulin AB (green). Phalloidin labels striated muscles in the umbrella wall (red), while nuclear DAPI is blue. (**a**) Each giant motor axon runs across the entire umbrella from the margin to the apex. In the margin, it divides into a few short branches (arrows) upon entering the nerve ring that circumvents the velum. (**b**) Higher magnification shows that α-tubulin AB label numerous thin filaments inside a giant axon, which represent long α-tubulin threads inside the giant syncytium. (**c**) Along the entire length of motor giants, there are more or less evenly spread lateral neural branches (arrows) running perpendicular to the giants and in parallel to the striated muscle fibers. (**d**) These lateral projections look like individual branches of the motor giants even at high magnification (arrows). However, the work by Weber et al. (1982) identified them as separate lateral neurons. (**e**) Lateral neurons branch extensively in the striated muscle layer. (**f**) Lateral neurons from neighboring motor giants (1 and 2) have overlapping innervation fields and even connect to each other. Abbreviations: *MGA* - motor giant axon; *INR* - inner nerve ring; *ONR* - outer nerve ring. Scale bars: a - 50 µm; b, d - 25 µm; c, e, f - 100 µm.

Individual motor giant axons, which run over the muscle tails of the subumbrella myoepithelial cells, were significantly protruding on the surface and visible in SEM images (Fig. 3c). The thickness of the motor giant axons could reach 35-40 µm at their base near the nerve ring that encircles the umbrella margin (Fig. 7a). They were significantly narrower in the upper part of the umbrella, in the range of 5-15 µm.

Near the nerve ring, each motor giant produced 3-4 short and thick branches that were entering the inner nerve ring (Fig. 7a). The nerve ring is the source that drives the activity of eight motor giant axons during both slow and escape/fast swimming in *Aglantha* (Roberts and Mackie, 1980; Meech and Mackie, 1993a; 1995).

Each motor giant axon also projected what looked like evenly spread, at 150-300 µm intervals, lateral branches perpendicular to its longitudinal orientation along with the circumferential striated muscles (Fig. 7c,d). These lateral branches have been identified by Weber et al. (1982) via dye-filling experiments as separate lateral neurons. The lateral neurons produced numerous thin neurites innervating and covering the entire body in the same focal plane where the circumferential striated muscles of the subumbrella were located (Fig. 7e). The thin branches from neighboring lateral neurons were overlapping and sometimes connecting each other (Fig. 7f). Giant motor axons, directly and via a population of lateral motor neurons, activate striated muscle cells of the subumbrella myoepithelium (Kerfoot et al., 1985).

Eight motor giant axons run in parallel and next to the eight radial digestive canals, from the margin of the umbrella to its apex. We have identified a single large axon running along the wall of each radial canal, labeled by α-tubulin AB (Fig. 8). Usually, this axon was masked by the bright α-tubulin immunostaining of the motor giant axons. However, in some preparations, motor giant axons were slightly moved to the side of the radial canals revealing the radial canal axon. This axon was produced by a single tripolar neuron located in the central area of the umbrella (Fig. 8b,c). One branch of this tripolar neuron was smaller and shorter, while the other two were running in the opposite direction over a substantial distance. This axon is a new addition to the system of large axons in *Aglantha*.

**Figure 8.**
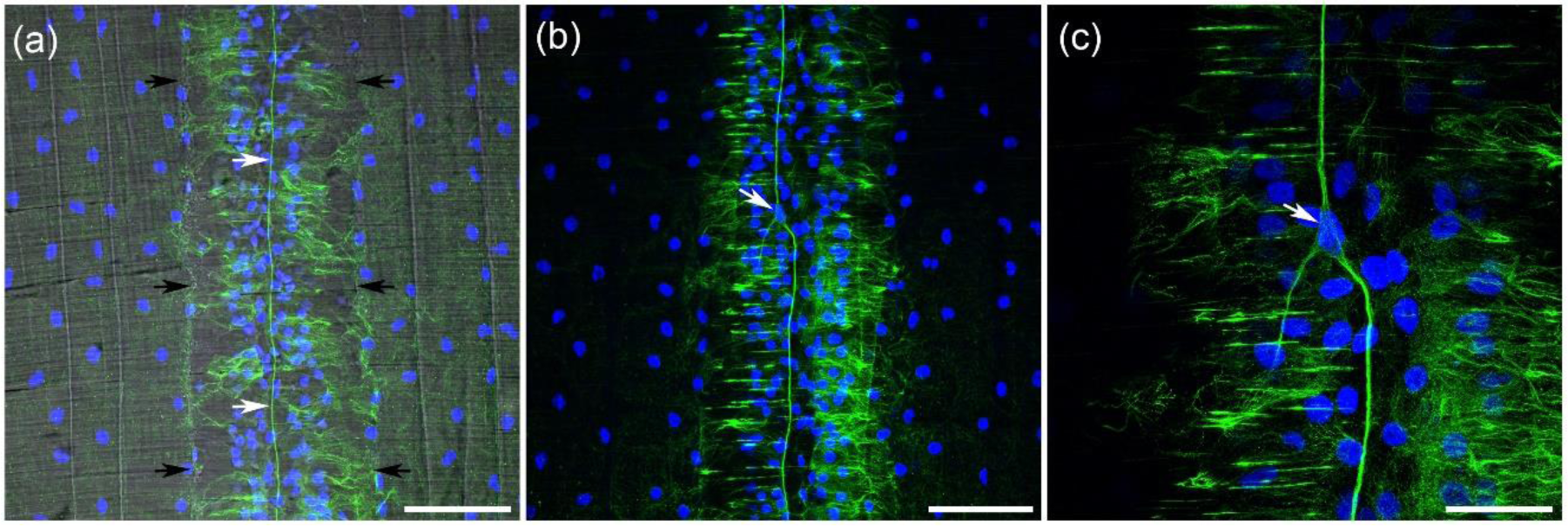
Large axon running along the radial digestive canal. α-Tubulin AB labeling is green, while nuclear DAPI is blue. (**a**) The radial digestive canal is outlined by black arrows in this image obtained with a transmitted light channel (TD). The axon (white arrows) in the middle of the canal is labeled by α-tubulin AB. (**b**) The long axon is produced by a single neuron (arrow). (**c**) This neuron (arrow) is tripolar with two larger branches running in opposite directions. Scale bars: a, b - 100 µm; c - 40 µm.

The nerve ring, which encircles the umbrella margin, is the central part of the nervous system in all hydrozoan jellyfishes (Fig. 9a; (Satterlie and Spencer, 1983; Satterlie, 2002)). In *Aglantha*, it consists of the inner nerve ring located closer to the subumbrella surface and the motor giant axons, and outer nerve ring with a ring giant axon located a little closer to the tentacles (Singla, 1978; Roberts and Mackie, 1980; Mackie and Meech, 1995b; a). Both the inner nerve ring and outer nerve ring were visible in α-tubulin immunolabeled preparations (Fig.9c). In addition to eight giant motor axons connecting to the nerve ring, it also gave rise to many thin neural processes forming a basal plexus around the ring (Fig. 9d).

**Figure 9.**
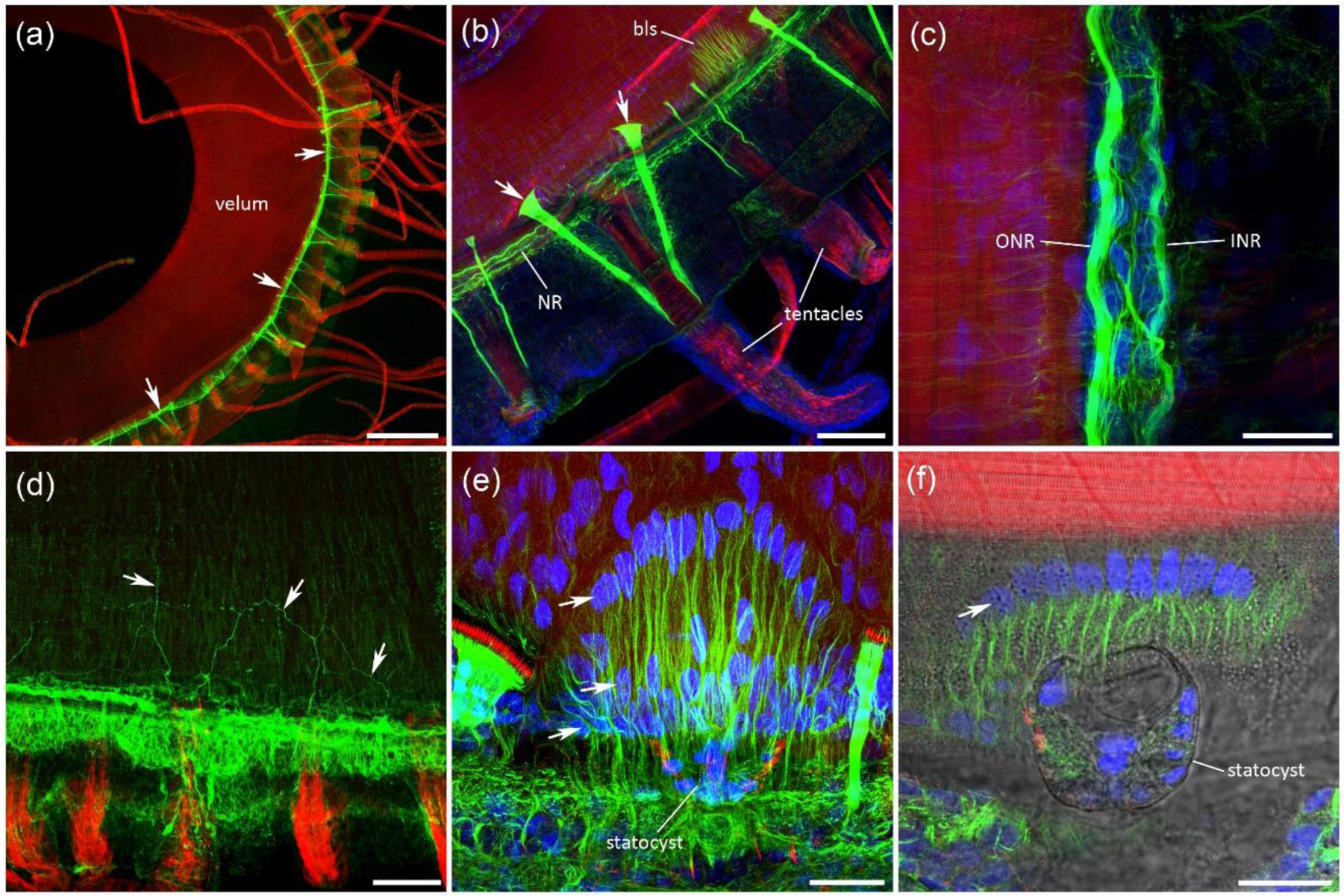
The nerve ring labeled by α-tubulin AB (green). Phalloidin is red; nuclear DAPI is blue. (**a**) The nerve ring circumvents the velum at the margin of the umbrella (arrows). (**b**) Crossing the nerve ring (*NR*) throughout its entire length, numerous short ‘cords’ are brightly labeled by α-tubulin AB - two on both sides of each tentacle base. Those cords are composed of sensory cells that form the comb pads on their wide side (arrows). (**c**) The nerve ring consists of two rings - outer nerve ring (*ONR*) and the inner nerve ring (*INR*), which can be seen at higher magnification. The outer nerve ring (*ONR*) contains a giant axon. (**d**) There are many thin processes originating from the ring that innervate the neighboring area of the body wall and form a basal plexus (arrows). (**e**) We have identified eight evenly spread bush-like structures attached to the nerve ring that consisted of multiple branches labeled by α-tubulin AB - also indicated as *bls* in (**b**). These structures were also characterized by several rows of tightly packed nuclei at different levels (arrows). (**f**) The addition of a transmitted light channel (TD) showed that at the base of each “bush” there was a statocyst. Scale bars: a - 500 µm; b, d - 100 µm; c, e, f - 25 µm.

### 3.4 Receptors - statocysts and comb pads

α-Tubulin immunostaining revealed two types of structures around the nerve ring in the umbrella margin area of *Aglantha*: 1) bush-like structures associated with statocysts, and 2) short and thick ‘cords’ crossing the nerve ring, which contained the ciliated sensory cells forming the comb pads (Fig. 9b).

There were eight bush-like structures evenly spread around the nerve ring. Each of them contained numerous α-tubulin immunoreactive branches radiating away from the base located right at the nerve ring level (Fig. 9b,e). The structure was flat and located in the epidermal layer. We do not know its functional role. However, the use of transmitted light (confocal differential interference contrast channel – TD) showed that there was a statocyst located at the base of each bush-like structure (Fig. 9f). The statocyst had a short row of sensory cilia on its side with each cilium surrounded by a ring of phalloidin-labeled microvilli. SEM confirmed the location of eight large statocysts, 30-50 µm in diameter, below the umbrella margin, next to the nerve ring location (Fig. 10a,b). The SEM also revealed a morphologically distinct area that corresponded to α-tubulin AB labeled bush-like structure attached to the statocyst (Fig. 10a). The statocysts have been shown to provide a sense of gravity and control direction of swimming in *Aglantha* (Mackie, 1980; Singla, 1983).

**Figure 10.**
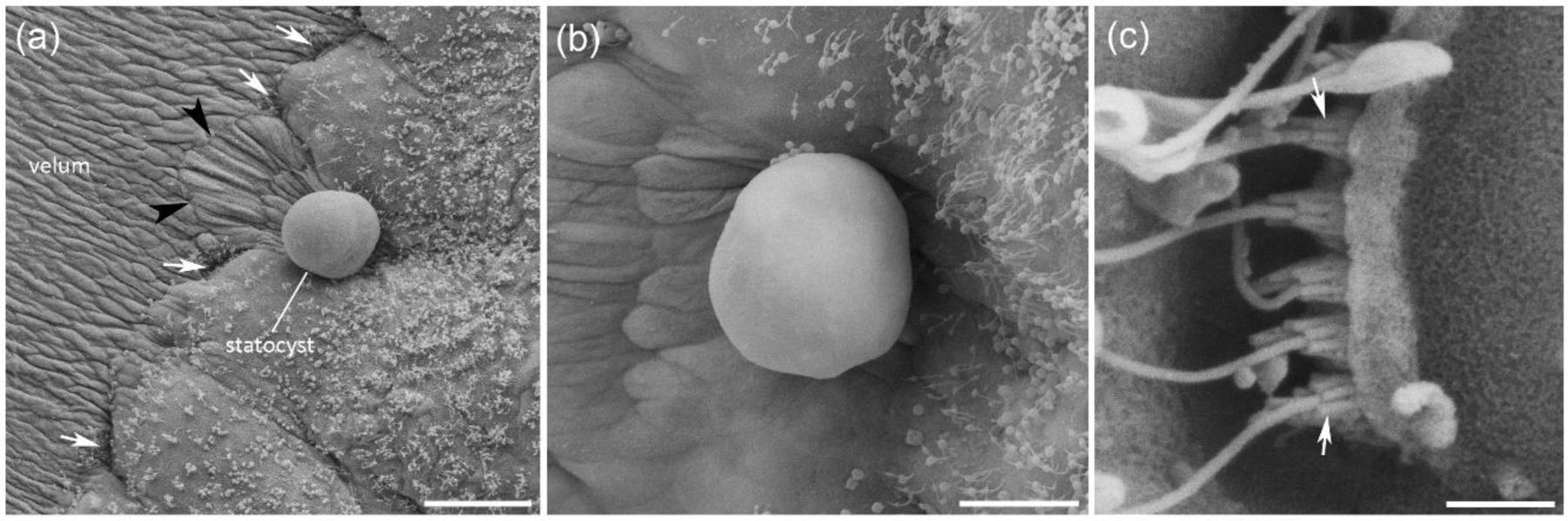
Scanning Electron Microscopy (SEM) of statocysts and comb pads. (**a**) The edge of the velum, right under the nerve ring, with a statocyst. White arrows point to the location of comb pads, while black arrowheads indicate the morphologically distinct area that corresponds to the previously described bush-like structure attached to the statocyst. (**b**) Statocyst. (**c**) Part of a comb pad at high magnification. Each long cilium in the comb pad is surrounded by a ring of microvilli (arrows), which have progressively differential length. Scale bars: a - 50 µm; b - 20 µm; c - 2 µm.

Comb pads are ciliated mechanosensory organs in *Aglantha* that presumably perceive disturbances in the water (Roberts and Mackie, 1980; Singla, 1983; Arkett et al., 1988). They are located along the line where the velum joins the umbrella margin (Fig. 10a). Each comb pad has several ciliated receptor cells. The smaller pads had 15-20 long sensory cilia, while the largest pad might have up to 50 cilia. Each long sensory cilium is surrounded at its base by a ring of short microvilli (Fig. 10c). There were 7 microvilli of different length in each ring, with the longest microvilli closer to the velum and shorter microvilli away from it (in agreement with Singla, 1983). α-tubulin AB or phalloidin did not label the long sensory cilia, which could be seen only with the transmitted light channel on the confocal microscope (Fig. 11c). However, the microvilli at the base of each cilium were labeled by phalloidin and could be seen as a row of brightly stained thorns or spikes, with each spike representing a ring of microvilli (Fig. 11b).

**Figure 11.**
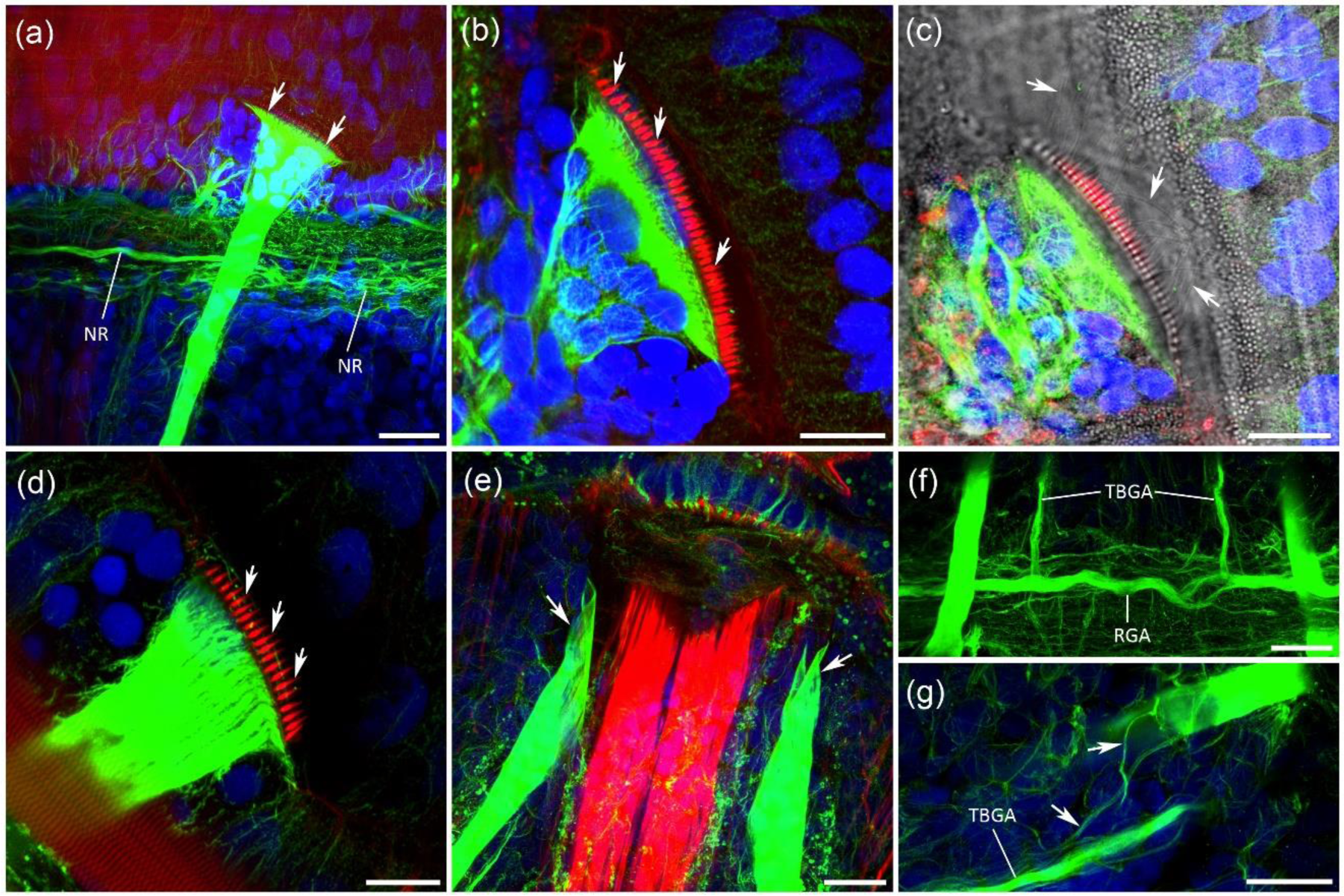
Sensory comb pads labeled by α-tubulin AB (green) and phalloidin (red), while nuclear DAPI is blue. (**a**) Comb pads are located on the wider end of the short and thick cords brightly labeled by α-tubulin AB. Those cords extend from the tentacle base, cross the nerve ring (*NR*) and widen into the actual comb pad at their end (arrows). (**b**) The microvilli surrounding the base of long comb pads cilia are brightly labeled by phalloidin. Each ring of microvilli looks like a short spike or thorn (arrows). (**c**) The long cilia themselves are not labeled by α-tubulin AB or phalloidin and could be seen only with a transmitted light channel (arrows). (**d**) Comb pads vary in size with smaller pads having only 15-20 sensory cilia with their microvilli labeled by phalloidin (arrows). (**e**) Comb pad cords (arrows) abruptly end on both sides of a muscle bundle connecting each tentacle base to the umbrella. (**f**) While crossing the nerve ring, comb pad cords run very close to the ring giant axon (*RGA*) almost making contact. The entire thickness of this optical section is only 5 µm. Note also that tentacle base giant axons (*TBGA*) contact the ring giant axon (*RGA*). (g) Short, thin branches (arrows) are connecting the tentacle base giant axons (*TBGA*) with the comb pad cords. Scale bars: a - 25 µm; b, c - 15 µm; d - 10 µm; e, f, g - 20 µm.

Each ciliated comb pad was located on the wider end of a short and thick ‘cord,’ brightly labeled by α-tubulin AB (Fig. 11a). These α-tubulin immunoreactive cords crossed the nerve ring close to the ring giant axon (Fig. 11f; video in Supplements), ran toward the tentacle base and ended on both sides of a bundle of muscles connecting tentacles to the umbrella margin (Fig. 9b, 11e, 12b, 15h). There were always two cords and two comb pads for each tentacle in *Aglantha* (Fig. 9a,b; 12b,c). Arkett et al. (1988) described comb pads as associated with the cell body of “enormous epithelial pad cell” with a “long axon-like process that runs toward the tentacle base.” We are not sure that each α-tubulin immunoreactive cord represents a single large epithelial cell (only serial reconstructions with transmission electron microscopy can answer that question). But, these ‘cords” are an integral part of the sensory comb pad structure, possibly acting as a secondary neural relay system. The cords end abruptly on both sides of muscle bundle at the base of each tentacle (Fig. 11e) and do not “fuse to become a large axon that runs along the tentacle” as was suggested by Arkett et al. (1988). Nevertheless, two giant axons at the tentacle base do produce thin short nerve branches that connect them to the pad cords (Fig. 11g).

### 3.5 Tentacles - innervation, cnidocytes, muscles and receptors

*Aglantha* has numerous tentacles attached to the low margin of its umbrella (Fig. 12a, 13a). Each tentacle has a distinct, short, and wide tentacle base connected to the margin and a long tentacle itself (Fig. 12b). Innervation of the *Aglantha* tentacles has been described morphologically and electrophysiologically with the emphasis on a giant tentacle axon, which runs through the entire length of a tentacle and is involved in triggering the escape response (Donaldson et al., 1980; Roberts and Mackie, 1980; Mackie et al., 1989; Bickell-Page and Mackie, 1991).

**Figure 12.**
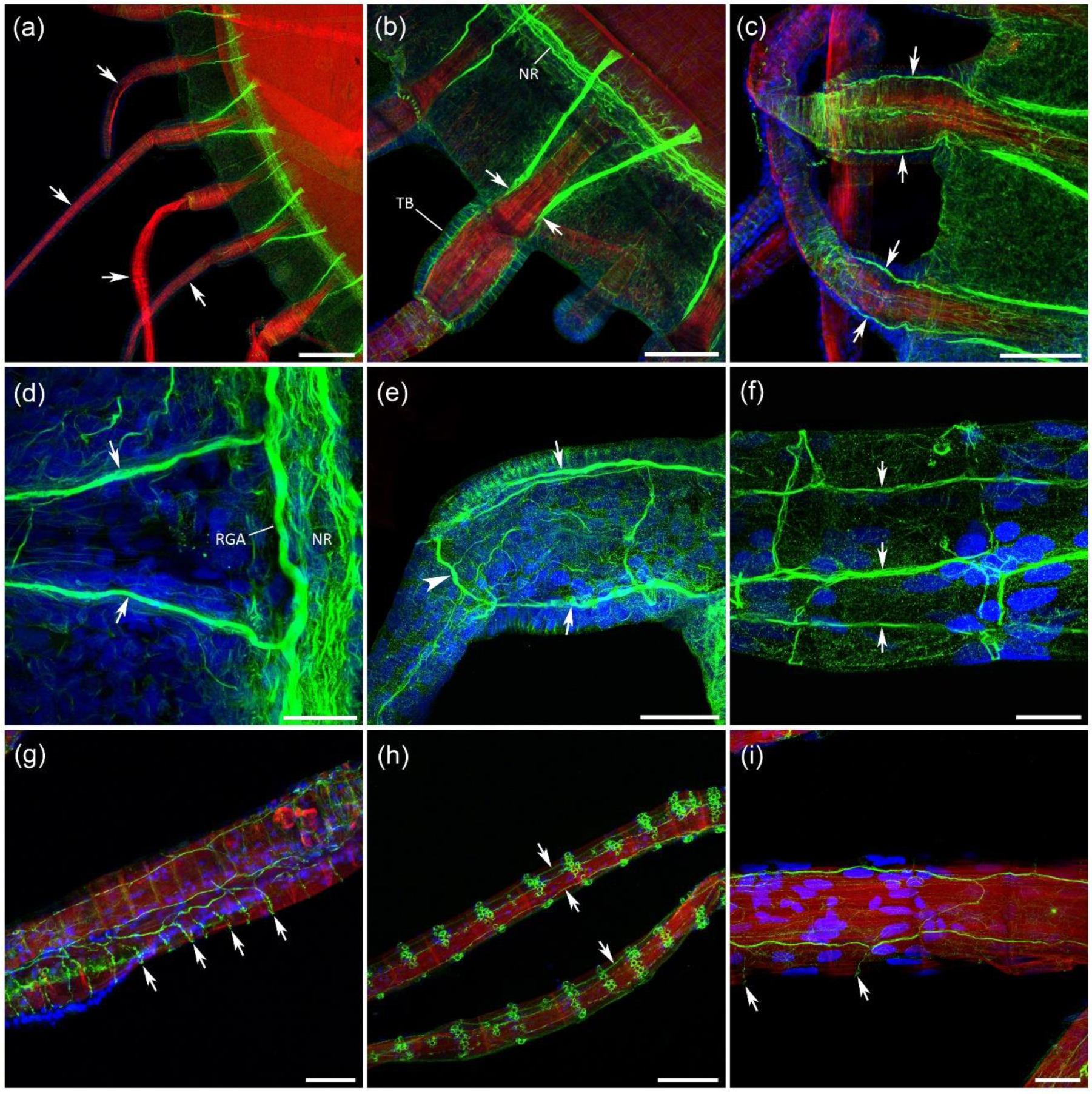
Innervation of the tentacles. α-Tubulin IR is green, phalloidin is red, while nuclear DAPI is blue. (**a**) Numerous thin and long tentacles (arrows) are attached to the margin of the umbrella in *Aglantha*. (**b**) Each tentacle has a distinct short tentacle base (*TB*). Two short cords labeled by α-tubulin AB, which form the sensory comb pads on one side, approach the base of each tentacle on the other side (arrows). (**c**) α-Tubulin AB also label two giant axons (arrows) that cross the base of each tentacle. (**d**) Two tentacle base giant axons (arrows) connect to the nerve ring (*NR*), making a visible contact with the ring giant axon (*RGA*). (**e**) The tentacle base giant axons (arrows) run across the base of the tentacle and form a ring-like connection to each other (arrowhead). (**f**) The ring, formed by tentacle base giant axons, gives rise to three axons (arrows) that run through the entire length of each tentacle. One axon (in the middle) is thicker than the other two and might correspond to the tentacle giant axon. (**g**) Close to their base, three axons produce a grid-like structure of connecting neural branches (arrows). (**h**) Further down, three axons (arrows) are observed crossing the entire length of the tentacles. (**i**) Fewer connecting branches (arrows) in the middle section of a tentacle. Scale bars: a - 200 µm; b, c, h - 100 µm; d - 30 µm; e - 40 µm; f - 20 µm; g - 50 µm; i - 25 µm.

**Figure 13.**
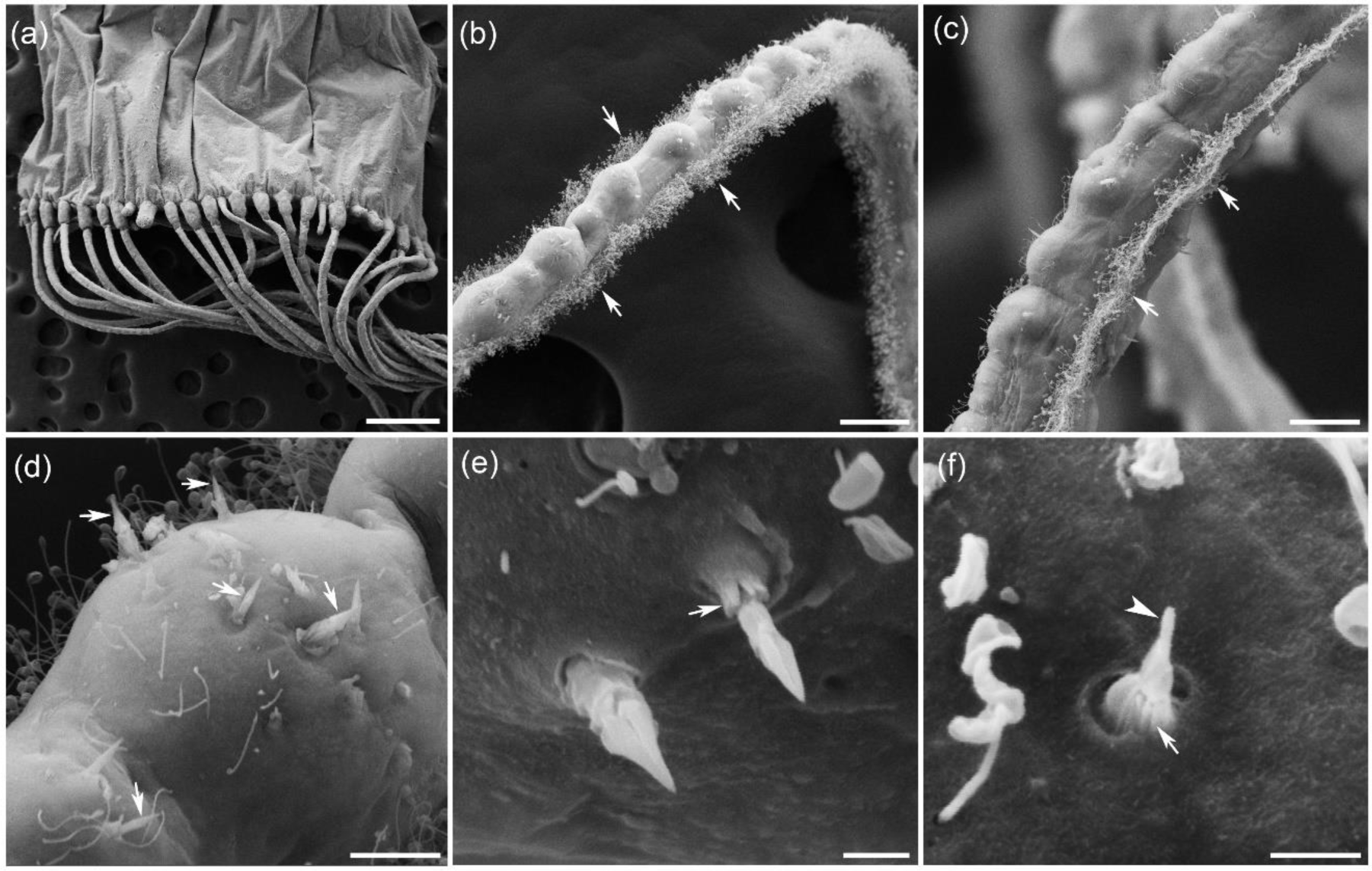
Scanning Electron Microscopy (SEM) of the tentacles. (**a**) Numerous tentacles originate at the margin of umbrella. (**b**, **c**) Each tentacle has a regularly spread spherical bulging areas along its length. There are two narrow bands of cilia (arrows) on both sides of each tentacle running through its entire length. (**d**) Bulging spheres contain cnidocytes (nematocytes). Some of the preparations had tentacles with many cnidocytes triggered and now demonstrating sharp protruding nematocysts seen on the surface of the tentacle (arrows). (**e**) Nematocysts are sharp, an arrow pointed and relatively short (two are shown here). Note the microvilli around the base of a nematocyst (arrow). (**f**) Some cnidocytes remained untriggered showing a single mechanosensory trigger cilium (cnidocil; indicated by arrowhead) on the top surrounded by a ring of microvilli (arrow) that tightly close the opening of cnidocyte. Scale bars: a - 500 µm; b - 50 µm; c - 25 µm; d - 10 µm; e, f - 2 µm.

Using anti-α-tubulin immunostaining, we have identified two giant axons, which ran from the nerve ring through the umbrella margin and the entire tentacle base on both sides (Fig. 12c). They corresponded to the proximal neurites of two giant tentacle neurons that gave rise to a single tentacle giant axon (Bickell-Page and Mackie, 1991). These two tentacle base giant axons established a direct connection with the ring giant axon from the outer nerve ring (Fig. 11f, 12d). It looked like the tentacle base axons merged with the ring giant (Fig. 11f, 12d). Previous electrophysiological experiments showed that action potentials in the tentacle giant axon had a one-to-one correspondence with the ring giant axon suggesting their electrical coupling (Donaldson et al., 1980; Roberts and Mackie, 1980). When the two tentacle base giant axons reach the top of the base, they connect to each other, forming a neural ring (Fig. 12e). Part of this neural ring could also be formed by independent ring neurons (Bickell-Page and Mackie, 1991).

The neural ring at the top of the tentacle base gave rise to three separate axons almost evenly spread around the ectodermal epithelium of the tentacle (Fig. 12f). One of the axons was thicker than the other two and may correspond to the previously described tentacle giant axon (Fig. 12f). All of the three axons were connected with short branches running perpendicular to the axons and circumferentially around the tentacle. Close to the tentacle base, there were very many such connectives, which created the grid-like appearance of the neural filaments (Fig. 12g). Further down to the tentacle tip, the number of connectives was significantly decreasing (Fig. 12i). All three axons could be followed through the entire length of each tentacle running in parallel to each other (Fig. 12h).

Tentacles in *Aglantha* are quite different from other hydrozoans. In addition to the system of giant axons, they also show two symmetrical bands of cilia on both sides of each tentacle. These cilia presumably assist in food collection by creating water currents along the tentacles (Mackie et al., 1989). The lateral cilia bands are seen on the SEM images (Fig. 13b,c).

Each tentacle had regularly spread spherical bulging areas along its length (Fig. 13b,c). All nematocytes (or cnidocytes) were located primarily on those bulging spheres (Fig. 13d). Some tentacles had nematocytes triggered before (or during) SEM fixation. The protruding nematocysts were relatively short, arrow-pointed, and sharp (Fig. 13e). There was a ring of small microvilli around the base of protruding nematocysts. ‘Not-triggered’ nematocytes had a short single sensory cilium (cnidocil) on the top with the ring of microvilli tightly closing around it (Fig. 13f).

α-Tubulin AB brightly labeled nematocytes on the tentacles. Clusters of nematocytes grouped around spherical bulging areas along the tentacles looked like colorful beads on the strand (Fig. 14a). There were very few nematocytes on the tentacles close to their base and umbrella margin, and much more near the tentacle tip (Fig. 14b,c). Each ‘not-triggered’ nematocyte had a cell body labeled by α-tubulin AB, one single large nucleus, a single mechanosensory cilium (cnidocil) also labeled by α-tubulin AB and a ring of small microvilli at the base of cnidocil, which were brightly labeled by phalloidin (Fig. 14d,e). We counted 7 microvilli around each sensory cilium in nematocyte (Fig. 14f). They were presumably closing the opening of the cnidocyte in the inactive, intact state.

**Figure 14.**
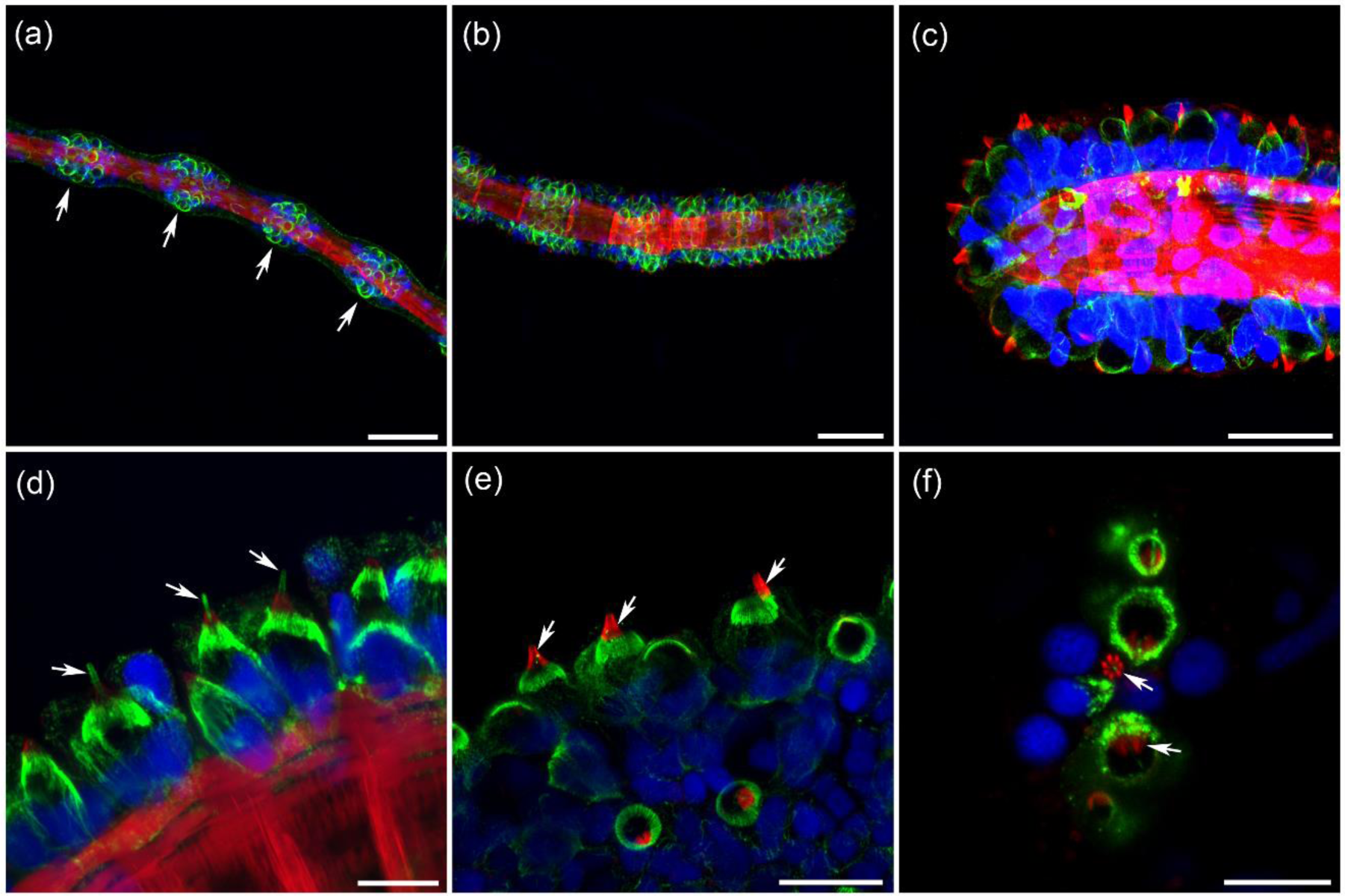
Nematocytes (cnidocytes) on the tentacles of *Aglantha*, labeled with α-tubulin AB (green), phalloidin (red) and nuclear DAPI (blue). (**a**) Nematocytes are labeled by α-tubulin AB (green). They are grouped in clusters (arrows) along the tentacles, which look like evenly spread spherical balls and remind of a strand of beads. (**b**, **c**) There are significantly more nematocytes at the tentacle end, and fewer closer to the tentacle base. Note the muscle bundles in the tentacles labeled by phalloidin (red). (**d**) Each intact cnidocyte is a cell with a single nucleus that has a mechanosensory single cilium (cnidocil) labeled by α-tubulin AB (arrows) on the top. (**e**, **f**) Phalloidin also labeled a ring of microvilli (arrows) at the base of the cnidocil cilium. In some preparations, we could count seven microvilli grouped around α-tubulin AB labeled cilium in the center (**f**). Scale bars: a, b - 50 µm; c - 25 µm; d, f - 10 µm; e - 20 µm.

Phalloidin labeled muscle fibers in the tentacles and the umbrella margin next to the tentacle base. Tentacles themselves had two groups of muscles: longitudinal muscle fibers and circular muscle bundles (Fig. 15a). From the tentacle base, ran a bundle of muscles through the umbrella margin up to the nerve ring (Fig. 15b,h). It looked like a root of the tentacle inside the umbrella. There were also muscle fibers between tentacle bases in the umbrella margin, which were apparently responsible for local movements and shape changes of the margin (Fig. 15b,c).

**Figure 15.**
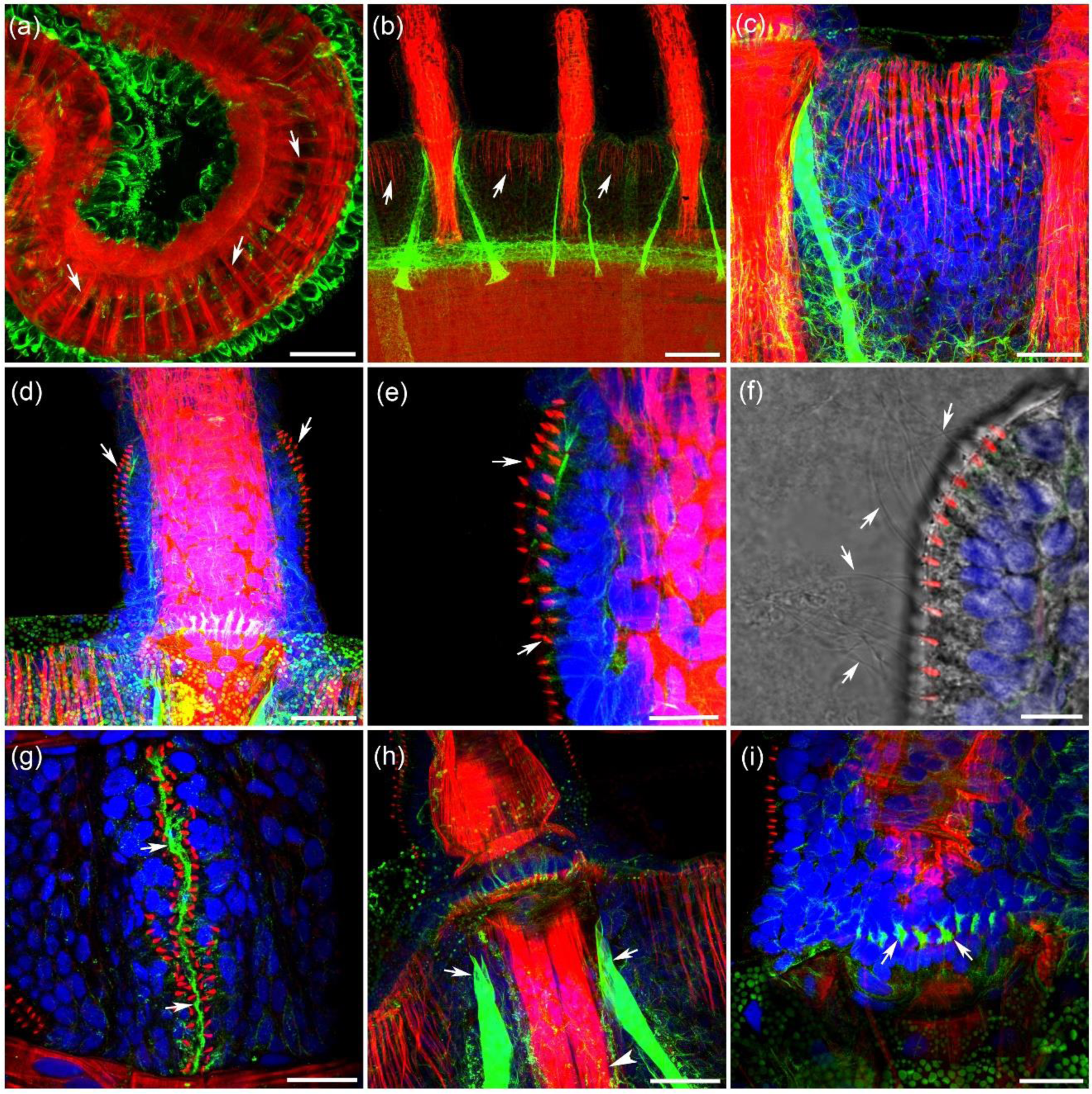
Muscles and ‘sensory’ cilia in the tentacles, labeled by phalloidin (red), α-tubulin AB (green) and nuclear DAPI (blue). (**a**) Tentacles are very muscular and intensively stained by phalloidin (red). There are two main groups of muscle fibers in each tentacle: circular muscle fibers (arrows) and long longitudinal muscle fibers. (**b**, **c**) Phalloidin also labels groups of muscle fibers at the very edge of the umbrella between the tentacles (arrows). (**d**, **e**) In addition, phalloidin brightly stained several rows of very short but pronounced ‘thorns’ or ‘spikes’ at the base of each tentacle (arrows). Each spike represented a compact group of short microvilli, which could be seen more clearly at SEM images (Figure 15). (**f**) The ring of these microvilli surrounded the long sensory cilia (arrows), which could be seen only with a transmitted light channel (TD) since they were not labeled by α-tubulin AB or phalloidin. (**g**) Neural connections to the sensory cilia at the tentacle base could be mediated by neurites (arrows) running along the cilia row and labeled by α-tubulin AB (green). (**h**) A powerful bundle of muscles (arrowhead) in the umbrella margin connects the base of each tentacle to deeper layers up to the ring nerve. Note the distant ends (arrows) of the sensory comb pads cords brightly labeled by α-tubulin AB. (**i**) A row of cell-like structures (arrows) is always labeled by α-tubulin AB at the tentacle base, where it connects to the umbrella margin. Scale bars: a, c, d, h - 40 µm; b - 100 µm; e - 15 µm; f - 10 µm; g, i - 25 µm.

Phalloidin also labeled several rows of short ‘spikes’ or ‘thorns’ running along the surface of each tentacle base (Fig. 15d,e). They looked very similar to the phalloidin-labeled rings of microvilli around long cilia in sensory comb pads described earlier. The use of the transmitted light channel revealed rows of long sensory cilia on the tentacle base, which were not labeled by α-tubulin antibody or phalloidin (Fig. 15f). Indeed, the phalloidin labeled ‘spikes’ or ‘thorns’ at the base of each long sensory cilia represented a ring of microvilli. It is important to note that there was always α-tubulin AB labeled neurite running along the rows of sensory cilia (and their microvilli), which could be involved in the relaying sensory information from them (Fig. 15g).

The rows of long sensory cilia could be seen during SEM imaging (Fig. 16b). At higher magnification, a ring of microvilli was seen surrounding the base of the sensory cilium (Fig. 16c). There were 7 microvilli of different length in each ring - very similar to the sensory cilia of the comb pads. These sensory cilia at the tentacle base have been studied before and identified as hair cells responsible for mechanoreception, which are capable of producing bursts of potentials in the nerve ring (Arkett et al., 1988). The umbrella margin also contained numerous cilia aggregated together in small groups, which probably served a sensory function (Fig. 16b).

**Figure 16.**
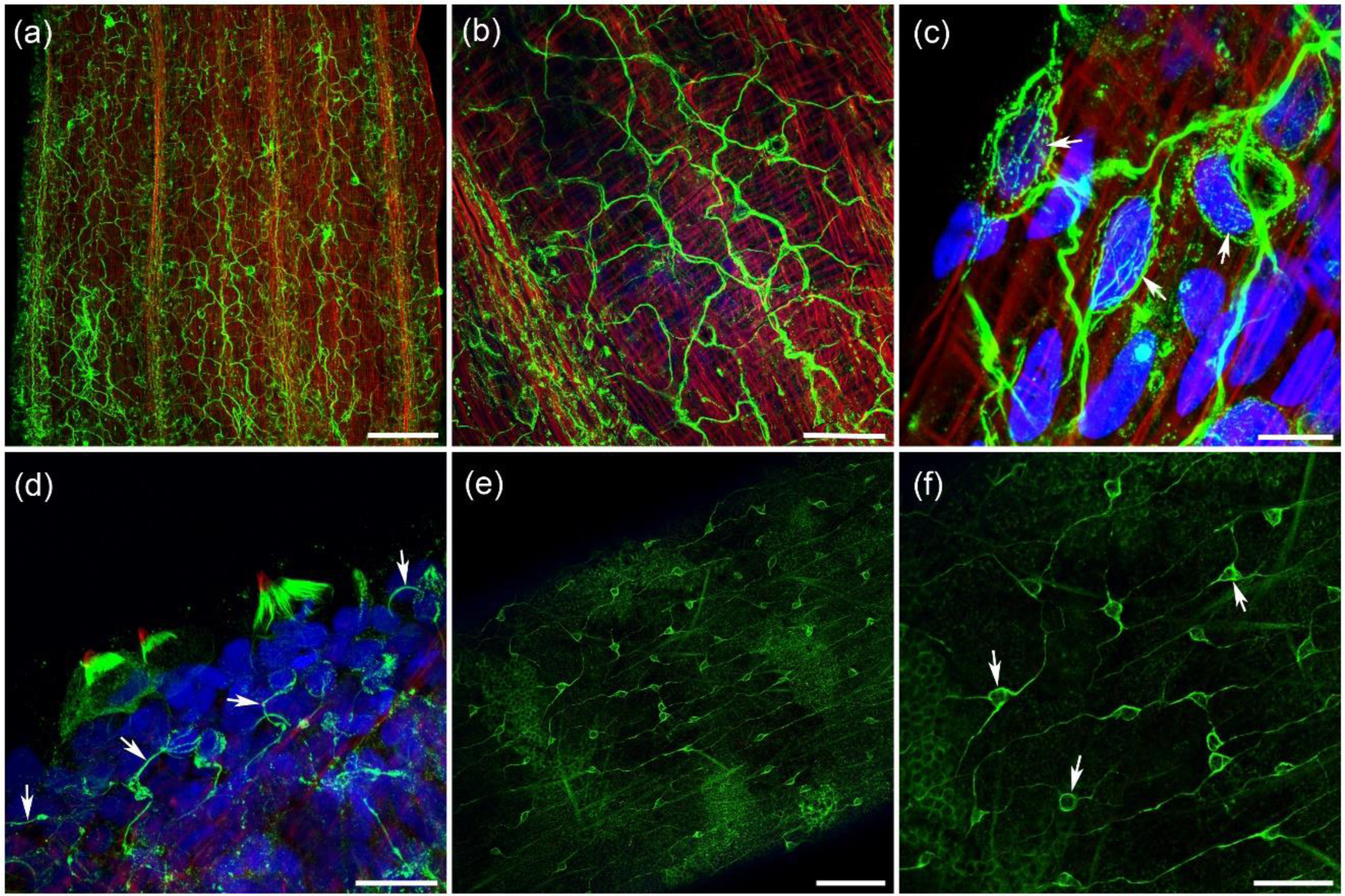
Scanning Electron Microscopy (SEM) of the base of *Aglantha* tentacles. (**a**) Each tentacle is attached to a short and wide base. They are seated on the narrow clearly defined disk on the outside of the velum, covered with numerous short cilia. (**b**) On the base of each tentacle, there are several rows of sensory cilia (arrows). (**c**) Each long sensory cilium is surrounded by a ring of microvilli of differential length (arrows). Scale bars: a - 200 µm; b - 50 µm; c - 1 µm.

### 3.6 FMRFa immunolabeling of additional neural elements

Anti-α-tubulin immunoreactivity proves to be a useful tool for identifying and visualizing neural elements in different animals. The question remains how comprehensive α-tubulin immunostaining is in identifying the majority of neural elements in cnidarians. To answer this question, we used α-tubulin antibody labeling together with another known marker of neural elements – (FM)RFamide-like immunoreactivity (RFa-IR – this abbreviation is used here to stress the fact that FMRFamide itself is not detected in *Aglantha* using RNA-seq). Antibody against FMRFamide is not very specific at identifying FMRFamide alone; they rather label a broad family of RFamide related peptides, most of which are markers for peptidergic neurons. Therefore, RFa-IR is consistently used to identify and study distinct neural populations in cnidaria (Grimmelikhuijzen and Spencer, 1984; Mackie et al., 1985; Satterlie, 2008; 2014; Satterlie and Eichinger, 2014).

The general distribution of RFa-IR in *Aglantha* in our experiments was similar to the description presented by Mackie et al. (1985). RFa-IR was primarily found in two areas: the umbrella margin, including the nerve ring and tentacles, and manubrium with lips around the mouth (Fig. 17a, 18a). The base of each tentacle contained a dense plexus of RFa-IR neurites on its aboral side (Fig. 17b). It also had a very distinct pair of symmetrically located RFa-IR multipolar neurons, which were called “star cells” by Mackie et al. (1985). These cells were located very close to the tentacle base giant axons and were <u>not</u> <u>labeled</u> by α-tubulin AB (Fig. 17b).

**Figure 17.**
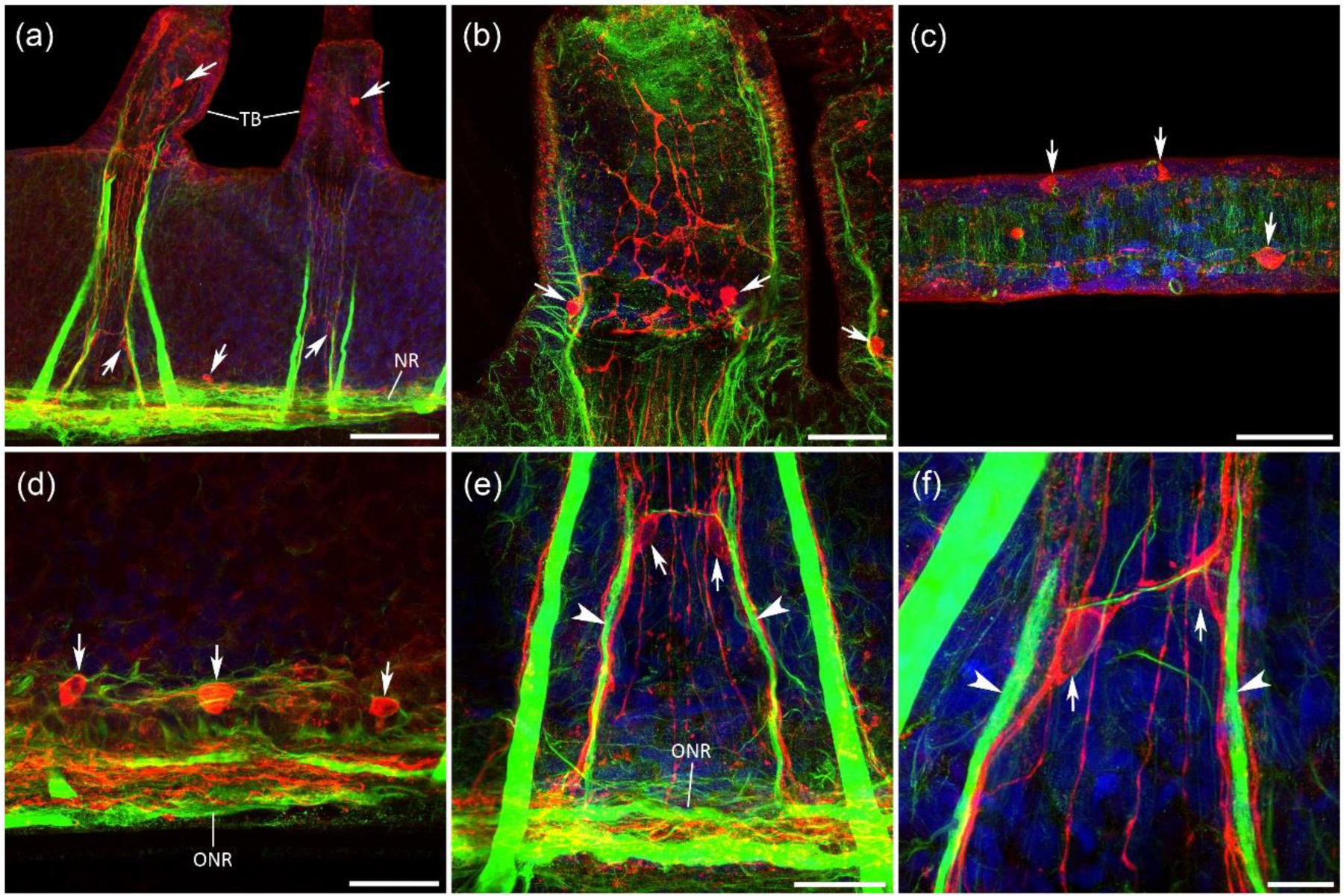
RFamide immunoreactivity (RFa-IR) in the umbrella margin and tentacles (labeled in red; while α-tubulin AB immunostaining is green). (**a**) Overview of immunoreactivity in the umbrella margin, from the nerve ring (*NR*) to the tentacle bases (*TB*). Arrows indicate some of the identified RFa-IR neurons. Note the tracts of RFamide-IR fibers running from the nerve ring to the tentacles. (**b**) Each tentacle base contains a plexus of RFa-immunoreactive processes and two larger neurons (arrows), which were identified as star cells by Mackie et al. (1985). (b) RFa immunoreactive cells in the tentacles (arrows). (**d**) Many round-shaped cells (arrows) are located along the outer nerve ring (*ONR*) through its entire length, along with numerous RFa immunoreactive processes inside the nerve ring. (**e**, **f**) A pair of large RFa immunoreactive neurons (arrows) has been identified next to each tentacle base giant axon (arrowheads) not far from where they enter the outer nerve ring (*ONR*). The thickest branches of these neurons (red) follow the tentacle base giant axons (green), while several smaller branches run in parallel contributing to the tract of IR fibers to the tentacles. Neurons in each pair are connected with a short connective. RFa immunoreactive neural processes (red) run along the thick comb pad cords (green). Note that RFa IR does not co-localize with α-tubulin IR, suggesting that α-tubulin AB do not label all neural elements in *Aglantha*. Scale bars: a - 100 µm; b, c - 40 µm; d, e - 30 µm; f - 10 µm.

**Figure 18.**
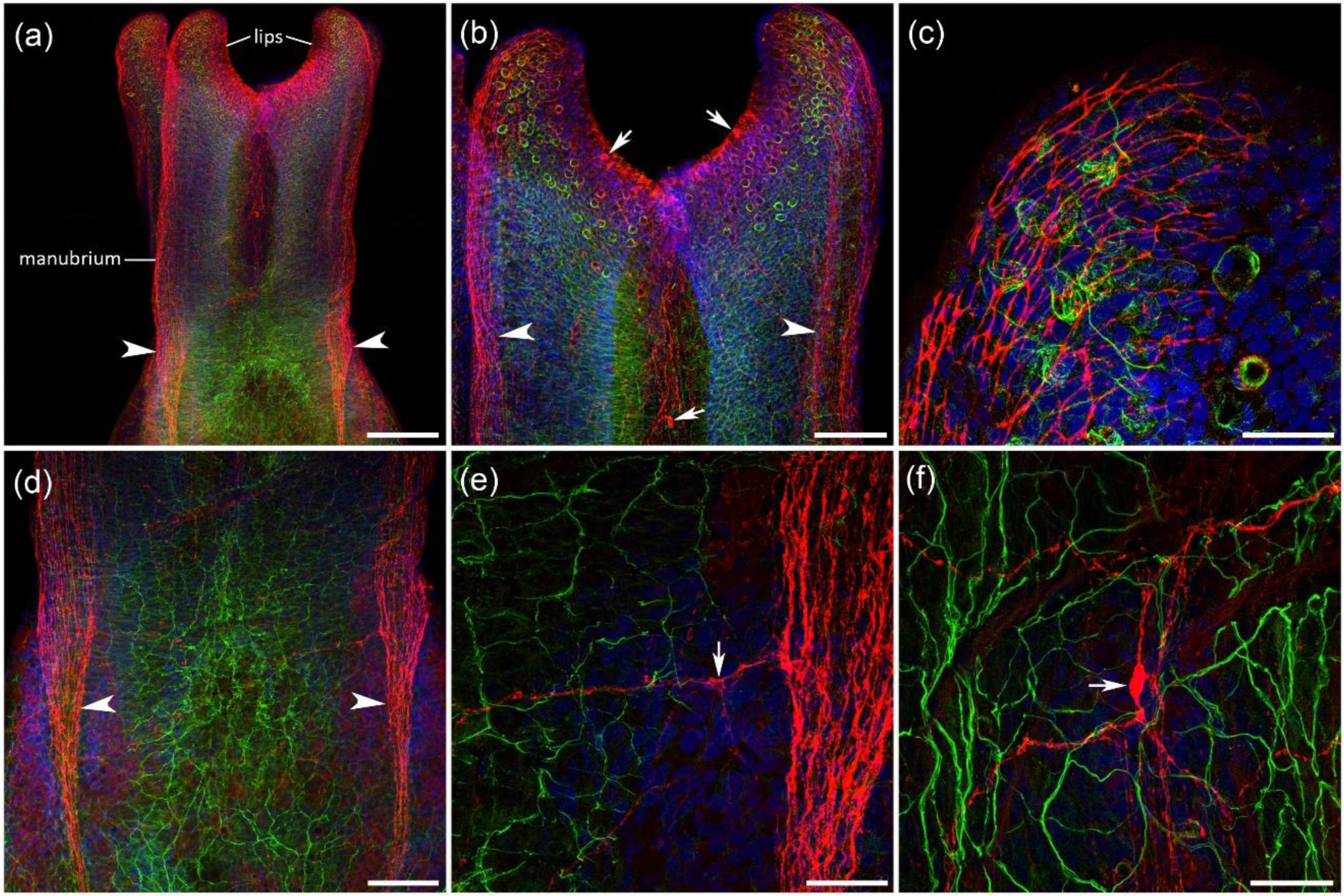
RFamide immunoreactivity in the manubrium and mouth (labeled in red; while α-tubulin AB immunostaining is green). (**a**) Manubrium and mouth with four lips. Arrowheads point to the RFa immunoreactive tracts, which run across the entire manubrium from peduncle to the tip of a lip. (**b**) Higher magnification of a mouth end of the manubrium. Arrows indicate some of the RFa immunoreactive neurons. (**c**) RFa immunoreactive plexus (red) at the tip of a lip. (**d**) The central part of the manubrium, where the narrow tracts expand into much wider stretches of RFa immunoreactive fibers (arrowheads). (**e**, **f**) RFa immunoreactive neurons (arrows) with their processes. Note that those neurons and their processes (red) do not overlap with α-tubulin immunoreactive plexus (green). Scale bars: a - 200 µm; b, d - 100 µm; c - 30 µm; e, f - 40 µm.

Tentacles themselves had numerous RFa-IR cell bodies along their entire length (Fig. 17c). Some of these cells could be sensory neurons (Mackie et al., 2003). RFa-IR sensory neurons were also identified along the outer nerve ring (Mackie et al., 1985; Mackie et al., 2003). The round-shaped labeled cell bodies were located regularly around the entire perimeter of the nerve ring (Fig. 17d). There were also numerous RFa-IR processes in the outer ring itself. Those filaments and immunoreactive sensory cells were also not labeled by α-tubulin AB.

A brightly labeled tract of RFa-IR neurites ran from the tentacle base to the nerve ring (Fig. 17a). A significant part of the neuronal processes in this tract was produced by a pair of relatively large RFa-IR neurons (15 µm in diameter), which we identified next to the tentacle base giant axons, not far from where they entered the outer nerve ring (Fig. 17e,f; video in Supplements). The cell bodies of these newly identified neurons were symmetrically located on each of the two tentacle base giant axons. A few larger axons produced by these RFa-IR neurons were attached to the tentacle base giant axons and followed them to the outer nerve ring in one direction and the tentacles in the other direction (Fig. 17e,f; video in Supplements). Several other thinner axons from these neurons contributed to a tract of RFa-IR neurites, running in the space between two tentacle base giant axons toward both the nerve ring and the tentacles. Two RFa-IR neurons were also connected by a short connective branch; The α-tubulin AB did not label those neurons.

Four distinct tracts of RFa-IR neurites crossed the manubrium from peduncle to the tip of four lips around the mouth (Fig. 18a). These tracks were widening closer to the lips and covered their entire outer subepithelial area (Fig. 18b,c). The inner area of the lips contained RFa-IR cell bodies (Fig. 18b), which were presumably sensory cells (Mackie et al., 1985, 2003). There were also several individual RFa-IR neurons with extensive and long branches between the tracts in the wall of the manubrium (Fig. 18e,f). All of the RFa-IR neurites and neurons in the manubrium did not have co-localization with α-tubulin immunoreactivity (Fig. 18c,e,f).

In summary, it appears that the majority, if not all neural elements, that are labeled by FMRFamide AB (antibody) are different from those revealed by α-tubulin immunostaining making these two neuronal markers complementary to each other in identifying the diversity of neural elements in *Aglantha*.

## 4. DISCUSSION

Neural systems in cnidarians and medusozoans, in particular, are very complex and diverse with multiple adaptive modifications reflecting long evolutionary histories in these lineages. Cnidarian neurons cannot be viewed as simple neural nets (see reviews (Satterlie and Spencer, 1983; Satterlie, 2002; Mackie, 2004; Satterlie, 2011; 2015b; a)), rather they are comparable in their heterogeneity and multifunctionality to bilaterian nervous systems.

At least 14 functional conductive systems have been physiologically identified in *Aglantha* including endodermal and ectodermal epithelial pathways (Mackie and Meech, 1995b; a; Mackie and Meech, 2000; Mackie et al., 2003) - the amazing example of neural complexity within a ‘simple” jellyfish! This makes *Aglantha* one of the best electrophysiologically characterized jellyfishes. Our work expanded the morphological bases of the neural organization of this important reference species for comparative and evolutionary neuroscience (Striedter et al., 2014).

### *Aglantha* as a highly specialized hydrozoan

According to the critical analyses performed by G. Mackie (2004), apart from more ancestral pacemaker systems nearly all “neural sub-systems <u>are unique to *Aglantha*</u>, or have been modified to the extent that their origins can no longer be recognized.” Furthermore, “the pacemaker system has undergone a drastic modification of its input-output relationships. Its primary output is no longer to the swim muscles (which have <u>lost the ability for myoid</u> <u>conduction</u>) but rather to neural components in the slow swimming motor pathway”(Mackie, 2004). Thus, we should view *Aglantha* as an example of what cnidarian nervous systems are ‘capable of achieving’ with distinct adaptations to mid-water pelagic lifestyle.

The concentration of about 800 small axons in two nerve rings of *Aglantha* (an outer and an inner nerve rings), together with associated neurons, forms so-called the **annulus** central nervous system (CNS). Two rings are physically connected, and, therefore, acting as a single centralized unit adapted to radial symmetry bodyplans (Mackie, 2004). The adaptive benefit of such annulus CNS is that both pacemakers and synaptic interactions are replicated at numerous points around the ring, providing both the redundancy of integrative systems and supporting the symmetry of the jellyfish. Such annulus central nerve rings can be analogous of the circumesophageal central nervous systems in bilaterians (Nielsen, 2012) also formed around the mouth and gut and, perhaps, for the same reasons.

The most exceptional feature of *Aglantha*, supporting its fast swimming and feeding behaviors, is the independent development of three types of giant axons (motor giant, ring giant, and tentacle giant axons). These axonal pathways do not have homologs in other hydrozoans studied so far. The system of giant axons in siphonophores (colonial hydrozoans) also evolved independently (Mackie, 1973; Mackie, 1978) from the *Aglantha* lineage.

The origins of giant axons occurred in parallel with the independent origin of striated muscles in the lineage led to *Aglantha*. However, the developmental and evolutionary source(s) of such origins are unclear. Giant axons are synthitiums with many nuclei, and could be formed either as a result of fusion events or incomplete cell divisions. Mills et al. suggested that giant axons might be evolved from excitable epithelium rather than from nerves – a clear example of the convergent evolution (Mills et al., 1985). This notice is supported by the fact that both the ring and tentacle giant axons are quite unusual in their ultrastructure too – their cytoplasm contains a large central vacuole running their entire length (Roberts and Mackie, 1980; Mackie et al., 1989). The motor giant axons are also physiologically unique in their ability to conduct two sorts of action potential (Na^+^ and Ca^2+^ spikes), which was never observed in bilaterians (Mackie and Meech, 1985). Motor giant axons and lateral neurons activate striated muscles in the umbrella and cause propulsion movement (Mackie, 2004).

By confirming and further illustrating many of the previous findings, we identified new neuronal elements in *Aglantha*. These are: 1) large axons along the radial digestive canals; 2) large RFa-IR cells located next to the tentacle base giant axons; 3) the system of three axons running along each tentacle (instead of just one tentacle giant axon described previously); 4) comb pad cords labeled by α-tubulin AB as a part of the neural system (it might be not a single epithelial cell as described by (Arkett et al., 1988)).

We have also clarified a few earlier observations. For example, we showed that tentacle base giant axons have a direct connection with the ring giant axon. Tentacle giant axon and ring giant axon were shown to be electrically coupled as they typically fired one-to-one (Roberts and Mackie, 1980; Mackie, 2004), but there were doubts that they contact histologically (Bickell-Page and Mackie, 1991). Now we show that there is indeed a direct connection between them. We have also found direct neural connections between comb pad cords and tentacle base giant neurons, which provide a morphological pathway between sensory comb pads and the neural system of the tentacles and central ring.

We suggest that three tentacle axons might have different functions, even though they are connected via short branches. The previously identified single tentacle giant axon (Mackie, 2004) is involved in a powerful tentacle retraction during the fast escape response. If the functional coupling between all three tentacle axons is high, then they would function as one unit during the escape. However, if their functional coupling is reduced (or they are uncoupled – e.g., by some modulators), the two smaller axons can be involved in smaller contractions during other behaviors.

In summary, the nervous system of *Aglantha* is composed of **seven** major components distributed across distinct regions of the body:

i. **The annulus CNS in the margin**. It consists of the inner nerve ring and outer nerve rings with the ring giant axon. The ring giant axon, as well as numerous other axons and neurites in the nerve ring, are labeled by α-tubulin AB. However, we found additional neurons and processes in the nerve ring that are labeled by FMRFa AB only. There are about 100-150 RFa IR cells around the nerve ring, including sensory-type neurons and large interneurons, as well as tracts of RFa IR neurites in the margin and tentacle base.
ii. **The system of motor giant neurons in the subumbrella** (total ∼500 neurons). This system includes eight motor giant axons running along the digestive canals and about 50 lateral neurons associated with each motor giant axon. All these neurons innervate striated muscles of the subumbrella myoepithelium. The lateral neurons and motor giant axons are labeled by α-tubulin AB. FMRFa AB didn’t label any of them. We didn’t detect any RFa IR neurons in the umbrella region above the nerve ring and up to the peduncle.
iii. **Large axons along radial digestive canals.** We have identified eight large axons running throughout the length of each of the eight radial digestive canals. They were produced by large tripolar neurons labeled by α-tubulin AB. They can control the functioning of the digestive canals. In fact, there were no neurons (labeled by α-tubulin AB or with RFa IR) other than digestive canal neurons, lateral neurons and motor giant axons in the subumbrella region between digestive canals, unlike in scyphomedusae (Satterlie and Eichinger, 2014), where neuronal elements where described in large subumbrella areas between the digestive canals. This further suggests a substantial centralization of the neural system in *Aglantha*. In contrast to ctenophores (Norekian and Moroz, 2019b; c; Norekian and Moroz, 2019a), we did not detect neuronal elements in the mesoglea of *Aglantha* using α-tubulin (or FMRFa AB).
iv. **Neurons in gonads**. We identified a diffusely distributed network of bipolar and multipolar neurons in gonads using α-tubulin AB, but there was no RFa IR in gonads. We could not morphologically link the nerve net in gonads to the rest of the nervous system, suggesting mainly local autonomous functions of this plexus.
v. **Neurons in the manubrium.** The entire manubrium is covered by a dense neural plexus labeled by α-tubulin AB. This neural network includes medium-sized neurons with their total number standing probably at about 200-300 cells. In addition, FMRFa AB labeled neural tracts, sensory cells in the lips, and individually standing neurons in the manubrium - totally about 50-100 cells, which were not stained by α-tubulin AB. There is electrophysiological evidence of coupling of this neural net with the annulus nervous system, which might include both neuronal and epithelial pathways (Mackie, 2004).
vi. **Tentacle nervous system.** We have identified three axons, running along the entire length of each tentacle, which were linked with several connective branches between them – all labeled by α-tubulin AB. The largest of the three was the previously described tentacle giant axon, instrumental in tentacle contraction during the escape. The α-tubulin AB also labeled two tentacle base giant axons, which may correspond to the proximal neurites of two giant tentacle neurons that gave rise to a single tentacle giant axon (Bickell-Page and Mackie, 1991). Interestingly, we did not see any α-tubulin AB labeled neuronal cell bodies in the tentacles (except nematocysts and nuclei associated with axons). However, there were at least 30-40 RFa IR neurons in each tentacle (an order of about 2000 neurons in all tentacles), including sensory-like cells, two “star cells” (Mackie et al., 1985) in the tentacle base and two newly identified neurons next to the tentacle base giant axons near the nerve ring.
vii. **Sensory systems** (see also above). First and foremost, it includes statocysts with their distributed components - for example, a newly identified and labeled by α-tubulin AB bush-like structure on the nerve ring.

*Aglantha* may lack photoreception and ocelli; show no ‘shadow response’ and lack visual neural circuitry, in contrast to other hydromedusae that live closer to the surface (Spencer and Arkett, 1984; Arkett and Spencer, 1986; Spencer, 1986). But, *Aglantha*’s very high vibrational sensitivity functionally acts as a trigger for the stereotyped fast escape adaptive behavior, which evolved to minimize a potential injury and predation from a diversity of cohabitants in mid-waters.

There are several types of ciliated, morphologically distinct receptors in *Aglantha* (counting nematocytes and putative chemoreceptors): (a) hair-type cell receptors on the base of the tentacles; (b) comb pads, which are connected to the rest of the nervous system via a α-tubulin AB labeled comb pad ‘cords’; (c) ciliated (mechano)receptors on the umbrella margin; (d) nematocysts; and (e) chemoreceptors.

Mouth with four lips and tentacles are two regions with high concentrations of nematocysts, which, as in other cnidarians, are considered to be sensory-effector parts of neural systems, possibly sharing the common origin with neurons (Marlow et al., 2012). We described anti-α-tubulin IR neurites along the rows of sensory cilia at the tentacle base and next to nematocytes presumably involved in their innervation.

Chemoreception has not been studied in *Aglantha*. However, the morphology of putative nitric oxide synthase (NOS) containing neurons and processes has been described in *Aglantha* with prominent labeling of putative chemosensory and other neurons in tentacles (Moroz et al., 2004). By comparing those and our recent datasets of α-tubulin-IR and RFa-IR cells, it looks like only a small fraction of putative (chemo)sensory cells in tentacles might have a colocalization with RFa-IR (putative neuropeptides) and NADPH-diaphorase labeling (a marker for NOS), while there is no detectable colocalization of α-tubulin-IR with other neuronal markers used here. These data further imply enormous molecular (possibly genetic) diversity of neuronal populations in *Aglantha* and a need for further neuroanatomical identification of neural circuits in this clade of metazoans.

At the broad scale, the employed markers visualize at least two distinct neural subsystems in *Aglantha* with the distinct structural and molecular organization of their composed neurons and processes. The α-tubulin AB mainly stained the nerves, giant axons, lateral motoneurons and neuronal processes. It corresponds more to our understanding of motor (effector) systems, which require prominent (elaborate) cytoskeleton organization. There are relatively few cell bodies labeled by α-tubulin AB. However, there are still more neurons labeled by tubulin AB in the umbrella than by FMRFa AB. The second subsystem includes sensory and interneurons (totaling ∼ 2500 cells in *Aglantha*), which could be mainly peptidergic and RFa IR.

In addition to the neural and sensory systems, another fluorescent marker phalloidin allowed us to identify at least eight types of different muscles in *Aglantha*, elaborating the lineage-specific effector innovations supporting complex behaviors. These are: 1) Striated muscles in subumbrella, which are part of myoepithelium; 2) Smooth spindle-shaped muscle fibers in the umbrella; 3) Thin longitudinal muscle fibers in the peduncle; 4) Muscle fibers in the manubrium wall, both longitudinal and circular, which create a grid-like layer; 5) Muscles in the margin of the umbrella between tentacles; 6) Longitudinal muscles in the tentacles; 7) Circular muscles in the tentacles; 8) Thick bundle of muscles that connects the tentacle base to the umbrella margin. Distinct molecular identities of these striated and smooth muscle cells are subjects to future investigations.

### *Aglantha* vs. other medusozoans

The hydromedusae are different significantly, both functionally and structurally, from the jellyfishes of two other Cnidaria classes - Scyphozoa and Cubozoa (Satterlie and Spencer, 1983; Satterlie and Nolen, 2001; Satterlie, 2002; Mackie, 2004; Satterlie, 2008; 2011; 2014; 2015b; a; Leitz, 2016). Each of this lineage shows examples of centralized neural systems (Mackie, 2004; Satterlie, 2011). The jellyfishes’ CNSs co-exist together with the diffuse nerve nets, which might represent a secondary adaptation to control two-dimensional myoepithelial subumbrella, and therefore, mediate synchronous contractions of the bell and jet-propulsion locomotion (Satterlie, 2015b; a).

Key functional differences in Hydrozoa impacting the overall organization of their integrative systems is the extensive development of gap junctions (Spencer and Satterlie, 1980; Mackie et al., 1984; Satterlie, 2002) and diversity of innexin genes (Moroz and Kohn, 2016). This supports both coupling of myoepithelial and neuronal cells required for neuromuscular coordination during feeding, swimming, and defense reactions (Satterlie, 2002; Mackie, 2004). Both scyphozoans and cubozoans explored different strategies to achieve synchronous controls of multiple muscles during swimming including the recruitment of different neural nets and chemical synaptic transmission vs. electric synapses (Satterlie, 2015a).

Specifically, Scyphozoans have developed two nerve nets in the subumbrella. The first is a motor (effector) highly polarized network, which is predominantly labeled by the anti-α-tubulin antibody. The second network has less polarization with multiple synaptic varicosities and mostly diffuse in its organization, which is predominantly labeled by (FM)RFamide AB (Satterlie and Eichinger, 2014). This later RFa IR network is thought to be modulatory. The rhopalia contain pacemakers that mediate the central control. They are analogs of bilaterian ganglia, although they have no or little connections between each other. In contrast, in Cubozoa, there is no diffuse neural RF-a IR net in the subumbrella, and the motor control is delegated to more developed rhopalia, which are connected by RFa IR (peptidergic) terminals to provide modulatory and sensory inputs as observed in the hydrozoan *Aglantha*.

Therefore, we conclude that in Cnidaria the centralization of neural systems might evolve at least three times in each of the medozozoan lineage (Scyphozoa, Cubozoa and Hydrozoa), and it could be associated with the sensory inputs and the integration of motor control as in *Aglantha* or cubozoans. The parallel neural centralization in Hydrozoa/*Aglantha* utilized the compression of multiple conductive systems, two nets at the junction of the subumbrella and velum, and elaborate electrical coupling.

## Conclusions

This paper provides a comprehensive atlas of the neuro-muscular organization in the hydrozoan *Aglantha digitale* with dozens of distinct neuronal, receptor and muscle cell types as a reference platform for future functional and comparative analyses.

First, even though *Aglantha* has been thoroughly investigated morphologically and physiologically in the past years and here, there are still many questions remain. For example, it is currently unclear how exactly multiple non-neuronal ectodermal, sensory, and neuronal pathways interact at the cellular level, and what diversity of synaptic and non-synaptic transmission exist. We also do not know the scope of signal molecules involved in these integrative neuronal mechanisms with hints that both nitric oxide (Moroz et al., 2004) and some unidentified peptides control feeding and locomotion. All these are topics for further investigation of this amazing biological system.

Second, we did not detect co-localization of α-tubulin IR with RFa IR. We also did not detect RFa-IR motor neurons further confirming a possibility that at least two structurally and molecularly distinct neural subsystems are present in *Aglantha* as in other jellyfishes (Satterlie, 2011; Eichinger and Satterlie, 2014; Satterlie and Eichinger, 2014). It would be safe to say that more novel neuronal elements and circuits would be revealed in *Aglantha* with additional cell-specific molecular markers as new signal molecules and transmitters would be identified using tools of single-cell genomics (Moroz, 2018).

Finally, three different bodyplans (in Scyphozoa, Cubozoa and Hydrozoa, respectively) and parallel evolution under the preexisting constraints (e.g., presence or absence of gap junctions or radial symmetry) led to the similar modes of locomotion by different means. All these three lineage-specific strategies also lead to three different forms of the CNS and, equally importantly, different diffuse nerve nets organizations, which reflect different recruitments of tubulin/cytoskeleton/polarization and peptides/signal molecules. Thus, we view the annulus-type CNS and rhopalia/ganglia as convergent centralization events. Similarly, some diffuse nerve nets can also be viewed as both ancient (evolutionary conservative) and secondarily specialized (derived) traits among Cnidaria lineages (and bilaterians – e.g., see (Moroz et al., 1997; Moroz, 2009)). *Aglantha* is an example of ‘the extreme case’ of neuromuscular adaptations driven by the prominent neuronal centralization in the form of the annulus nervous system to support the fast escape swimming and elaborate behavior repertoire.

## Data Availability Statement

The data that support the findings of this study are available from the corresponding author upon request.

## Acknowledgments

We thank FHL for their excellent facilities including the Nikon Laser Scanning confocal microscope and Scanning Electron Microscope. This work was supported by the National Science Foundation (grants 1146575, 1557923, 1548121 and 1645219).

## Role of the authors

All authors had full access to all the data in the study and take responsibility for the integrity of the data and the accuracy of the data analysis. TPN and LLM share authorship equally. Research design: TPN, LLM. Acquisition of data: TPN, LLM. Analysis and interpretation of data: TPN, LLM. Drafting of the article: TPN, LLM. Funding: LLM.

## LITERATURE CITED

1. Arcila D, Orti G, Vari R, Armbruster JW, Stiassny MLJ, Ko KD, Sabaj MH, Lundberg J, Revell LJ, Betancur RR. 2017. Genome-wide interrogation advances resolution of recalcitrant groups in the tree of life. Nat Ecol Evol 1(2):20.

2. Arkett S, A., Mackie GO, Meech RW. 1988. Hair cell mechanoreception in the jellyfish *Aglantha digitale*. J Exp Biol 135:329–342.

3. Arkett SA, Spencer AN. 1986. Neuronal mechanisms of a hydromedusan shadow reflex. I. Identified reflex components and sequence of events. J Comp Physiol 159:201–213.

4. Bickell-Page LR, Mackie GO. 1991. Tentacle autotomy in the hydromedusa *Aglantha digitale* (Cnidaria): an ultrastructural and neurophysiological analysis. . Phil Trans R Soc Lond B 331:155–170.

5. Brusca RC, Brusca GJ. 2003. Invertebrates. Sunderland, Massachusetts: Sinauer Associates, Inc. 936 p.

6. Bullock TH, Horridge GA. 1965. Structure and Function in the Nervous Systems of Invertebrates. San Francisco: Freeman.

7. Donaldson S, Mackie GO, Roberts A. 1980. Preliminary observations on escape swimming and giant neurons in *Aglantha digitale* (Hydromedusae: Trachylina). Can J Zool 58:549–552.

8. Eichinger JM, Satterlie RA. 2014. Organization of the ectodermal nervous structures in medusae: cubomedusae. Biol Bull 226(1):41–55.

9. Greenberg MJ, Payza K, Nachman RJ, Holman GM, DA. P. 1988. Relationship between the FMRFamide-related peptides and other peptide families. Peptides 9:125–135.

10. Grimmelikhuijzen CJP, Spencer AN. 1984. FMRFamide immunoreactivity in the nervous system of the medusa *Polyorchis penicillatus*. J Comp Neurology 230(3):361–371.

11. Kerfoot PA, Mackie GO, Meech RW, Roberts A, Singla CL. 1985. Neuromuscular transmission in the jellyfish *Aglantha digitale*. J Exp Biol 116:1–25.

12. Kozloff EN. 1990. Invertebrates. Philadelphia: Sounders College Publishing. 866 p.

13. Leitz T. 2016. Cnidaria. In: Schmidt-Rhaesa A, Harzsch S, Purschke G, eds. Structure and Evolution of Invertebrate Nervous Systems. Oxford: Oxford University Press. p 26–47.

14. Mackie G, Meech R. 1995a. Central circuitry in the jellyfish *Aglantha*. I: The relay system. J Exp Biol 198(Pt 11):2261–2270.

15. Mackie G, Meech R. 1995b. Central circuitry in the jellyfish Aglantha. II: The ring giant and carrier systems. J Exp Biol 198(Pt 11):2271–2278.

16. Mackie GO. 1973. Report on giant nerve fibres in *Nanomia*. Publ Seto Mar Lab 20:745–756.

17. Mackie GO. 1978. Coordination in physonectid siphonophores. Mar Behav Physiol 5:325–346.

18. Mackie GO. 1980. Slow swimming and cyclical ‘fishing’ behavior in *Aglantha digitale* (Hydromedusae: Trachylina). Can J Fish Aquat Sci 37:1550–1556.

19. Mackie GO. 1990. The elementary nervous sytems revisited. American Zoologist 30:907–920.

20. Mackie GO. 2004. Central neural circuitry in the jellyfish *Aglantha*: a model ’simple nervous system’. Neurosignals 13(1-2):5–19.

21. Mackie GO, Anderson PAV, Singla CL. 1984. Apparent absence of gap junctions in two classes of Cnidaria. Biol Bull 167:120–123.

22. Mackie GO, Marx RM, Meech RW. 2003. Central circuitry in the jellyfish *Aglantha digitale* IV. Pathways coordinating feeding behaviour. J Exp Biol 206(Pt 14):2487–2505.

23. Mackie GO, Meech RW. 1985. Separate sodium and calcium spikes in the same axon. Nature 313(6005):791–793.

24. Mackie GO, Meech RW. 2000. Central circuitry in the jellyfish *Aglantha digitale*. III. The rootlet and pacemaker systems. J Exp Biol 203(Pt 12):1797–1807.

25. Mackie GO, Meech RW. 2008. Nerves in the endodermal canals of hydromedusae and their role in swimming inhibition. Invert Neurosci 8(4):199–209.

26. Mackie GO, Nielsen C, Singla CL. 1989. The tentacle cilia of *Aglantha digitale* (Hydrozoa: Trachylina) and their control. Acta Zool (Stockh) 70:133–141.

27. Mackie GO, Singla CL, Stell WK. 1985. Distribution of nerve elements showing FMRFamide-like immunoreactivity in hydromedusae. Acta Zool (Stockholm) 66:199–210.

28. Marlow H, Roettinger E, Boekhout M, Martindale MQ. 2012. Functional roles of Notch signaling in the cnidarian *Nematostella vectensis*. Dev Biol 362(2):295–308.

29. Meech RW. 2015. Electrogenesis in the lower Metazoa and implications for neuronal integration. J Exp Biol 218(Pt 4):537–550.

30. Meech RW, Mackie GO. 1993a. Ionic currents in giant motor axons of the jellyfish, *Aglantha digitale*. J Neurophysiol 69(3):884–893.

31. Meech RW, Mackie GO. 1993b. Potassium channel family in giant motor axons of *Aglantha digitale*. J Neurophysiol 69(3):894–901.

32. Meech RW, Mackie GO. 1995. Synaptic potentials and threshold currents underlying spike production in motor giant axons of *Aglantha digitale*. J Neurophysiol 74(4):1662–1670.

33. Mills CE, Mackie GO, Singla CL. 1985. Giant nerve axons and escape swimming in *Amphogona apicata* with notes on other hydromedusae. Can J Zool 63:2221–2224.

34. Moroz LL. 2009. On the independent origins of complex brains and neurons. Brain, behavior and evolution 74(3):177–190.

35. Moroz LL. 2014. The genealogy of genealogy of neurons. Commun Integr Biol 7(6):e993269.

36. Moroz LL. 2015. Biodiversity Meets Neuroscience: From the Sequencing Ship (Ship-Seq) to Deciphering Parallel Evolution of Neural Systems in Omic’s Era. Integr Comp Biol 55(6):1005–1017.

37. Moroz LL. 2018. NeuroSystematics and Periodic System of Neurons: Model vs Reference Species at Single-cell Resolution. ACS Chem Neurosci 9(8):1884–1903.

38. Moroz LL, Kocot KM, Citarella MR, Dosung S, Norekian TP, Povolotskaya IS, Grigorenko AP, Dailey C, Berezikov E, Buckley KM, Ptitsyn A, Reshetov D, Mukherjee K, Moroz TP, Bobkova Y, Yu F, Kapitonov VV, Jurka J, Bobkov YV, Swore JJ, Girardo DO, Fodor A, Gusev F, Sanford R, Bruders R, Kittler E, Mills CE, Rast JP, Derelle R, Solovyev VV, Kondrashov FA, Swalla BJ, Sweedler JV, Rogaev EI, Halanych KM, Kohn AB. 2014. The ctenophore genome and the evolutionary origins of neural systems. Nature 510(7503):109–114.

39. Moroz LL, Kohn AB. 2016. Independent origins of neurons and synapses: insights from ctenophores. Philos Trans R Soc Lond B Biol Sci 371(1685):20150041.

40. Moroz LL, Meech RW, Sweedler JV, Mackie GO. 2004. Nitric oxide regulates swimming in the jellyfish *Aglantha digitale*. J Comp Neurol 471(1):26–36.

41. Moroz LL, Sudlow LC, Jing J, Gillette R. 1997. Serotonin-immunoreactivity in peripheral tissues of the opisthobranch molluscs *Pleurobranchaea californica* and *Tritonia diomedea*. J Comp Neurol 382(2):176–188.

42. Nielsen C. 2012. Animal Evolution: Interrelationships of the living phyla. Oxford: Oxford University Press.

43. Norekian TP, Moroz LL. 2016. Development of neuromuscular organization in the ctenophore *Pleurobrachia bachei*. J Comp Neurol 524(1):136–151.

44. Norekian TP, Moroz LL. 2019a. Comparative neuroanatomy of ctenophores: Neural and muscular systems in Euplokamis dunlapae and related species. J Comp Neurol:in press.

45. Norekian TP, Moroz LL. 2019b. Neural System and Receptor Diversity in the ctenophore Beroe abyssicola. J Comparative Neurology:in review.

46. Norekian TP, Moroz LL. 2019c. Neuromuscular organization of the Ctenophore *Pleurobrachia bachei*. J Comp Neurol 527(2):406–436.

47. Roberts A, Mackie GO. 1980. The giant axon escape system of a hydrozoan medusa, *Aglantha digitale*. J Exp Biol 84:303–318.

48. Satterlie RA. 2002. Neural control of swimming in jellyfish: A comparative story. Canad J Zool 80:1654–1669.

49. Satterlie RA. 2008. Control of swimming in the hydrozoan jellyfish Aequorea victoria: subumbrellar organization and local inhibition. J Exp Biol 211(Pt 21):3467–3477.

50. Satterlie RA. 2011. Do jellyfish have central nervous systems? J Exp Biol 214(Pt 8):1215–1223.

51. Satterlie RA. 2014. Multiple conducting systems in the cubomedusa Carybdea marsupialis. Biol Bull 227(3):274–284.

52. Satterlie RA. 2015a. Cnidarian Nerve Nets and Neuromuscular Efficiency. Integr Comp Biol 55(6):1050–1057.

53. Satterlie RA. 2015b. The search for ancestral nervous systems: an integrative and comparative approach. J Exp Biol 218(Pt 4):612–617.

54. Satterlie RA, Eichinger JM. 2014. Organization of the ectodermal nervous structures in jellyfish: scyphomedusae. Biol Bull 226(1):29–40.

55. Satterlie RA, Nolen TG. 2001. Why do cubomedusae have only four swim pacemakers? J Exp Biol 204(Pt 8):1413–1419.

56. Satterlie RA, Spencer AN. 1983. Neuronal control of locomotion in hydrozoan medusae. J Comp Physiol A150:195–206.

57. Simion P, Philippe H, Baurain D, Jager M, Richter DJ, Di Franco A, Roure B, Satoh N, Queinnec E, Ereskovsky A, Lapebie P, Corre E, Delsuc F, King N, Worheide G, Manuel M. 2017. A Large and Consistent Phylogenomic Dataset Supports Sponges as the Sister Group to All Other Animals. Curr Biol 27(7):958–967.

58. Singla CL. 1978. Locomotion and neuromuscular system of *Aglantha digitale*. Cell Tissue Res 188(2):317–327.

59. Singla CL. 1983. Fine structure of the sensory receptors of *Aglantha digitale* (Hydromedusae: Trachylina). Cell Tissue Res 231(2):415–425.

60. Spencer AN. 1986. Neuronal mechanisms of a hydromedusan shadow reflex. II. Graded response of reflex components, possible mechanisms of photic integration and functional significance. J Comp Physiol 159:215–225.

61. Spencer AN, Arkett SA. 1984. Radial symmetry and the organization of central neurones in a hydrozoan jellyfish. J Exp Biol 110:69–90.

62. Spencer AN, Satterlie RA. 1980. Electrical and dye coupling in an identified group of neurons in a coelenterate. J Neurobiol 11(1):13–19.

63. Steinmetz PR, Kraus JE, Larroux C, Hammel JU, Amon-Hassenzahl A, Houliston E, Worheide G, Nickel M, Degnan BM, Technau U. 2012. Independent evolution of striated muscles in cnidarians and bilaterians. Nature 487(7406):231–234.

64. Striedter GF, Belgard TG, Chen CC, Davis FP, Finlay BL, Gunturkun O, Hale ME, Harris JA, Hecht EE, Hof PR, Hofmann HA, Holland LZ, Iwaniuk AN, Jarvis ED, Karten HJ, Katz PS, Kristan WB, Macagno ER, Mitra PP, Moroz LL, Preuss TM, Ragsdale CW, Sherwood CC, Stevens CF, Stuttgen MC, Tsumoto T, Wilczynski W. 2014. NSF workshop report: discovering general principles of nervous system organization by comparing brain maps across species. J Comp Neurol 522(7):1445–1453.

65. Weber C, Singla CL, Kerfoot PA. 1982. Microanatomy of the subumbrellar motor innervation in *Aglantha digitale* (Hydromedusae: Trachylina). Cell Tissue Res 223(2):305–312.

66. Wehland J, Willingham MC. 1983. A rat monoclonal antibody reacting specifically with the tyrosylated form of alpha-tubulin. II. Effects on cell movement, organization of microtubules, and intermediate filaments, and arrangement of Golgi elements. J Cell Biol 97(5 Pt 1):1476–1490.

67. Wehland J, Willingham MC, Sandoval IV. 1983. A rat monoclonal antibody reacting specifically with the tyrosylated form of alpha-tubulin. I. Biochemical characterization, effects on microtubule polymerization in vitro, and microtubule polymerization and organization in vivo. J Cell Biol 97(5 Pt 1):1467–1475.

68. Whelan NV, Kocot KM, Moroz LL, Halanych KM. 2015. Error, signal, and the placement of Ctenophora sister to all other animals. Proc Natl Acad Sci U S A 112(18):5773–5778.

69. Whelan NV, Kocot KM, Moroz TP, Mukherjee K, Williams P, Paulay G, Moroz LL, Halanych KM. 2017. Ctenophore relationships and their placement as the sister group to all other animals. Nat Ecol Evol 1(11):1737–1746.

70. Wulf E, Deboben A, Bautz FA, Faulstich H, Wieland T. 1979. Fluorescent phallotoxin, a tool for the visualization of cellular actin. Proc Natl Acad Sci U S A 76(9):4498–4502.

